# Decrypting the languages of allostery in membrane-bound K-Ras4B using four complementary *in silico* approaches

**DOI:** 10.1101/2023.07.14.549022

**Authors:** Matteo Castelli, Filippo Marchetti, Sílvia Osuna, A. Sofia F. Oliveira, Adrian J. Mulholland, Stefano A. Serapian, Giorgio Colombo

**Affiliations:** Department of Chemistry, University of Pavia, viale T. Taramelli 12, 27100 Pavia, ITALY; INSTM, via G. Giusti 9, 50121 Florence, ITALY; E4 Computer Engineering, via Martiri delle libertà 66, 42019 Scandiano (RE), ITALY; Institut de Química Computacional i Catàlisi (IQCC) and Departament de Química, Universitat de Girona, Girona, Catalonia E-17071, SPAIN; ICREA, Barcelona, Catalonia E-08010, SPAIN; Centre for Computational Chemistry, School of Chemistry, University of Bristol, Bristol BS8 1TS, U.K; School of Biochemistry, University of Bristol, Bristol BS8 1TD, U.K.

**Keywords:** K-Ras4B, allostery, shortest path map, distance fluctuation, molecular dynamics, D-NEMD

## Abstract

Proteins are evolved molecular machines whose diverse biochemical functions can be dynamically regulated by allostery, through which even faraway protein residues can conformationally communicate. Allostery can express itself in different ways, akin to different “languages”: allosteric control pathways predominating in an unperturbed protein are superseded by others as soon as a perturbation arises—*e.g.*, a mutation—that alters its function (pathologically or not). Accurately modeling these often-unintuitive phenomena could therefore help explain functional changes in a specific protein whenever they are unclear. Unbiased molecular dynamics (MD) simulations are a possibility; however, since allostery can operate at longer timescales than those accessible by MD, simulations require integration with a reliable method able to, *e.g.*, detect regions of incipient allosteric change or likely perturbation pathways. Several methods exist but are typically applied singularly: we argue their joint application, in a “multilingual” approach, could significantly enrich this picture. To prove this, we perform unbiased MD simulations (∼100 µs) of the widely studied, allosterically active oncotarget K-Ras4B, solvated and embedded in a phospholipid membrane, proceeding to decrypt its allostery using four showcase “languages”: *Distance Fluctuation* analysis and the *Shortest Path Map* capture allosteric communication hotspots at equilibrium; *Anisotropic Thermal Diffusion* and *Dynamical Non-Equilibrium MD* simulations assess them once the GTP that oncogenically activates K-Ras4B is, respectively, either superheated or hydrolyzed. “Languages” provide a uniquely articulate, mutually coherent, experimentally consistent picture of allostery in K-Ras4B. At equilibrium, pathways stretch from the membrane-embedded hypervariable region all the way to the active site, touching known flexible allosteric “switches” and proposed pockets. Upon GTP cleavage/perturbation, known to inactivate K-Ras4B, allosteric signals most reverberate on switches and interfaces that recruit effector proteins. Our work highlights the benefits of integrating different allostery detection methods with unbiased MD.

## Introduction

Proteins are efficient and versatile machines that support most biochemical processes in cells.^1,2^ To meet these requirements, proteins populate a diverse array of structures that are intrinsically dynamic^3-5^ and are required to sustain well defined and finely tuned functional motions.^6-8^

Allostery^6,9,10^ is a fundamental mechanism regulating such functions, whereby distal parts of a protein or multimeric complex (often not intuitively linkable to active sites or binding interfaces) dynamically communicate with each other, with far-reaching repercussions on cellular activities: examples of the many aspects inextricably dependent on allostery^10^ include enzyme function,^11-13^ protein folding by chaperones,^14-16^ signal transduction,^17^ and regulation of transcription and metabolism. Understanding of allostery has been constantly expanding since seminal studies in the 1960s^18^ and 1980s,^19^ with the topic being extensively reviewed and reformulated.^4,7-10,20-24^

Dynamic (residue-mediated) allosteric communication pathways arise even in the more conformationally constrained proteins,^6-8,19^ mediating the transition between distinct energy-wells in biomolecules characterized by well-defined “folding funnel” minima.^5,10,23,25^ Such minima associated with the native structure can be rugged and different sub-states can exist as rapidly interconverting conformational ensembles of slightly different functional/nonfunctional forms. Allostery is the key factor regulating transitions between these forms so that they coexist in precisely the right proportions required to ensure biological functions. In this framework, a targeted allosteric perturbation is what helps prompt a (physiological) change of function.^10,21,22^ Such perturbation is typically introduced by either one or more specific post-translational modifications (PTMs), or an endogenous ligand binding at an allosteric site, or complexation with another protein, or by cleavage of a substrate. As a result, conformational equilibria are subtly altered, ushering in a “population shift”,^4,7,8,10,20,21,23,25-27^ as allosteric signals are relayed for a new biochemical event to occur, often far from the perturbation site(s). These events can result, *e.g.*, in the modification of the properties of a particular interface, with consequent promotion or disruption of another protein’s recognition;^28,29^ facilitation of conformational change;^14^ substrate binding or release;^30^ the turnover rate of a molecular machine;^31^ and regulation of enzymatic reactivity,^13^ which is of particular interest in the field of biocatalysis.^8,11,12,26,30,32^

With such a delicate set of conformational equilibria required for normal biological functions, it is unsurprising that aberrant allosteric perturbations are enough to disrupt the physiological balance among different conformational populations and lead to a number of pathologies.^22^ Indeed, an emerging therapeutic strategy^21,25,29^ is to design small-molecule allosteric modulators^25,28,29^ that bind to and interfere with identified allosteric pockets, providing a possible alternative to ineffective or toxic *ortho*steric ligands.^21,25,28^

The signal-transducing GTPase K-Ras^36,37^ in its most abundant oncogenic isoform K-Ras4B^36^ (Figure 1) is a textbook case of a small but allosterically complex protein (residues 2-185 when mature), consisting of a globular catalytic G domain (residues 2-166; Figure 1; yellow, purple, black, and cyan) followed by a flexible hypervariable region (HVR; Figure 1a,b; salmon) terminating with a farnesylated Cys185 that is responsible for its incorporation into the cellular membrane (Figure 1a).^34,37,38^ Under healthy conditions, membrane-bound K-Ras cycles between an *active* (GTP-/Mg^2+^-bound) state (Figure 1a,b) and an *inactive* one wherein GTP has been hydrolyzed to GDP (Figure 1c).^35^ Only when K-Ras is active, key regions of its G domain (switches I and II; Figure 1)^37,39^ can be allosterically remodeled to recruit and help activate various effectors,^36,37,39^ which then trigger appropriate signaling cascades. Subsequent K-Ras deactivation through GTP hydrolysis also requires switches I and II to adopt specific conformations^37,39,40^ and it is greatly facilitated^39,40^ by the recruitment of a GTPase-activating protein (GAP) which immobilizes a catalytically crucial^40^ glutamine (Gln61) in K-Ras and administers an equally crucial^40^ arginine (Arg789 in GAP numbering; Figure S1b). Hydrolysis results in an inactive G domain with switches allosterically incapacitated to recruit effectors. In as many as 1 in 10 cancers,^41,42^ K-Ras4B is seen to undergo a plethora of different mutations^36,37,41^ and/or PTMs^37^ that, in ways that are not always allosterically clear^37,41^ (bar steric disruption of the K-Ras4B–GAP interface, or catalytic interference, *e.g.*, through mutation of key residues Lys16 or Gln61; Figure S1b and Figure 1),^40^ hinder GTP cleavage and thus trap the protein in a harmful *hyperactive* state. In fact, while clinically relevant mutations overwhelmingly^42^ concentrate in the P-loop and particularly on Gly12 and Gly13 (Figure 1; black), they can be found all over the GTPase.

**Figure 1.**
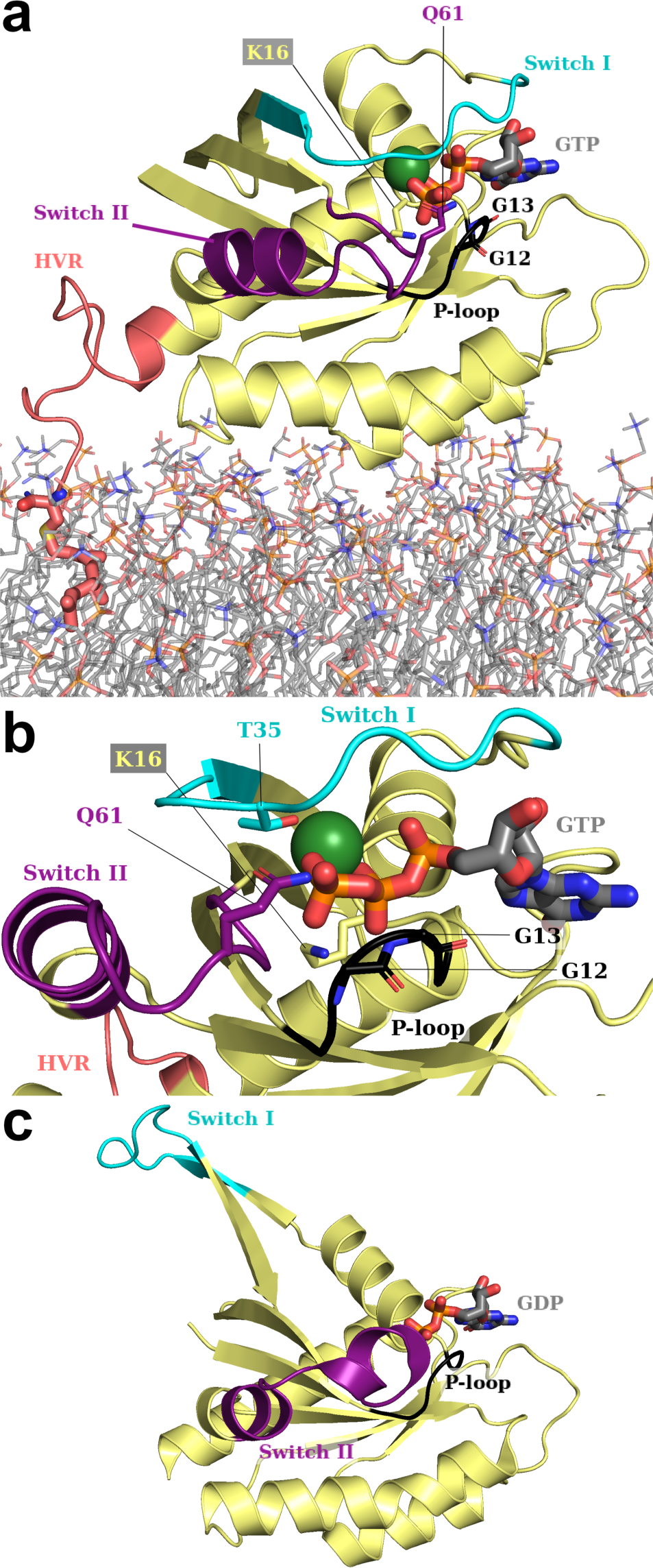
Structural overview of K-Ras4B, active (as simulated in this work) and inactive. (a) Zoom on K-Ras4B as it appears in our starting structure (Figure S1a). Modelled after PDB ID: 6vjj,^33^ the G domain (residues 2-166; yellow) is represented with its salient features (see main text): switches I and II (in cyan and purple, respectively); and the P-loop (black; in a ‘reduced’ definition). Catalytically relevant (Lys16, Gln61) and mutation-prone residues (Gly12, Gly13) are explicitly labeled and rendered as sticks, with C atoms and labels in the same color as their parent feature. The hypervariable region (HVR; residues 167-185; salmon) and the POPS/POPC phospholipid bilayer (lines) in which it is embedded through farnesylated Cys185 (C as large salmon spheres) are modeled from previous simulations.^34^ (b) Further zoom on K-Ras4B after a ∼90° clockwise rotation perpendicular to the plane of the phospholipid bilayer (which is here omitted), revealing the same features. (c) Inactive K-Ras4B after GTP hydrolysis to GDP (PDB ID: 6mqg),^35^ in the same orientation as (a) and showing the same features (HVR not present). In all panels, GTP (GDP in (c)) is represented as thicker sticks. H, Na^+^, Cl^-^ and solvent are omitted for clarity. Color code for explicit non-C atoms/ions: Mg^2+^ (absent from (c)): dark green; P: orange; O: red; N: blue; S: yellow.

In light of its relevance to oncology and unsuitability for orthosteric GTP-competitive inhibition, copious efforts have been made to identify allosteric propagation routes and hotspots in K-Ras and its mutants (*e.g.*, ^36,41,43,44^), culminating in the recent (titanic) experimental effort by Weng *et al.*^41^ to produce a comprehensive allosteric map of the effects of K-Ras mutations at all possible sequence positions. Significantly for this work, K-Ras4B dynamics and their variations in mutants have also been intensely studied from a computational point of view.^34,37,38,43,45-54^ For a complete picture, we certainly refer the reader to Pantsar’s comprehensive review^37^ but, for example, molecular dynamics (MD) simulations have confirmed: (i) that G domain switches are flexible^47,53^ and flexibility is altered in different activation states^47,53^ and oncogenic mutants;^50^ (ii) that the G domain can fold back on the HVR and is affected by it,^45^ even when K-Ras4B is embedded in the membrane;^34,46^ (iii) that this interaction is weakened in oncogenic mutants;^49^ (iv) that the GTPase can rotate when anchored to the membrane;^34^ and that (v) there exist several allosteric pockets in the G domain^43,53^ (later explicitly assessed in the aforementioned experimental allosteric map).^41^ Yet, despite all these findings and the clear medical need, covalent allosteric inhibitor Sotorasib (AMG510),^55^ which rescues GTP cleavage in the oft-recurring K-Ras4B mutant of Gly12 to Cys (G12C; *cf.* Gly12 in Figure 1), remains the only clinically approved option so far.^41^ The noncovalent pan-mutant inhibitor BI-2865, impeding the restoration of the GTP-active state by preventing nucleotide exchange, was reported as late as April 2023.^56^ Such paucity of medical options is indicative of the difficult druggability of K-Ras4B and the allosteric diversity of its oncogenic aberrations,^36,37,41^ justifying the continued interest in this GTPase and its allosteric mechanisms.

Computational techniques^21,23,57^ are essential to obtain an atomistic understanding of the determinants of allosteric regulation.^21,23,24^ While the timescales accessible by atomistic simulations (now on the microsecond scale) remain on the short side when it comes to capturing complete allosteric conformational changes,^27^ a number of solutions, including coarse-graining and enhanced sampling MD, have become available over the years (*cf.* ^21,23,27,58^), as well as the combination with machine learning.^59^ Markov State Model (MSM) analysis, Replica Exchange MD, and Principal Component Analysis (PCA) have also been applied specifically to K- Ras4B.^47,48,50,53^

In this context, there exist atomistic-resolution methods rooted in unbiased MD that, while starting from different initial assumptions, aim to tackle the question of unveiling the atomistic and mechanistic details of allosteric modulation and the key residues involved.

Here, exploiting the considerable trove of allosteric information on K-Ras4B, we prove that a synergistic combination of four such techniques can readily interrogate 5 µs long atomistic MD simulations of K-Ras4B to provide a unique level of insight into its allosteric regulation, with each technique addressing a different aspect. Our overarching aim is to extract consensus information that will point us to the key residues/substructures and essential dynamic traits mediating allosteric regulation in K-Ras (K-Ras4B). Chosen methods to decrypt allostery include: (i) residue-pair distance fluctuation (DF) analysis, normally used to detect allosteric “crosstalk” patterns in large proteins and complexes thereof (*e.g.*, ^14,16,60^); (ii) the shortest path map (SPM) method devised by Osuna and coworkers^12,30^ to suggest (even distal) mutations that are most likely to allosterically impact on an enzyme’s reactivity as desired; (iii) the dynamical nonequilibrium MD (D-NEMD) simulations initially developed by Ciccotti and Jacucci,^61^ which maps allosteric signal transduction from effector and/or substrate binding sites in receptors and enzymes;^17,62-66^ and (iv) anisotropic thermal diffusion (ATD)^67^ which entails heating an effector in its binding site within a (supercooled) protein or complex, and monitoring whereto the allosteric message propagates. Based on our results, we offer a view of how often seemingly independent approaches, developed from different perspectives, in reality speak allosteric languages that are not so unintelligible and can be employed cooperatively, uncovering more information than if used on their own. We propose the concerted use of these techniques to unveil consensus mechanistic determinants and lay the basis for a more complete molecular understanding of allosteric regulation.

## Computational Methods

### General Procedure

To begin with, atomistic molecular dynamics (MD) simulations of membrane-embedded K-Ras4B were set up and conducted, as discussed below, in 20 independent replicas. From these, we directly derived distance fluctuation (DF)^14,16^ matrices and shortest path map (SPM)^12,30^ as recounted later (*i.e.*, the first two out of the four allosteric languages considered in our study). In addition, a subset of frames isolated from these equilibrium MD simulations serve as the starting point for further nonequilibrium MD simulations *per* the final two allostery detection methods (languages), *i.e.*, dynamical nonequilibrium MD (D-NEMD)^17^ and anisotropic thermal diffusion (ATD).^67^

### System Setup

All equilibrium MD simulations were begun from a single representative structure (Figure S1a) of mature, active K-Ras4B embedded in a previously equilibrated^34^ phospholipid bilayer (23% POPS, 77% POPC) through its farnesylated Cys185 (henceforth Fcy185), and solvated in an aqueous solution of 0.1 M ionic strength—achieved through the presence of appropriate amounts of Na^+^ and Cl^-^ ions—extending on both sides of the bilayer. We prepared this structure as recounted below, ensuring that characteristics of mature K-Ras4B were modeled as closely as possible.^37^

Our template for modeling the G domain was a 1.40 Å-resolution crystal structure of wild-type (active) K-Ras4B (PDB ID: 6vjj; *cf.* description in Figure 1a,b).^33^ From this structure we proceeded with the *PyMol* package^68^ as follows: we removed the co-crystallized domain of effector RAF1, Cl^—^ ions, and small molecule-ligands; the non-hydrolyzable GTP mimic GMPPNP was converted into GTP through *in situ* replacement, by an oxygen, of the nitrogen atom connecting Pγ and Pβ; and two *N*-terminal residues were stripped to obtain the mature form of the G domain (plus Lys167), complete with an *N-*acetyl cap on Thr2. To reflect its catalytically active^40^ conformation, the Gln61 sidechain was rotated inwards to match its observed orientation in the presence of a GAP (Figure S1b; PDB ID: 1wq1);^69^ all other atoms and molecules, including crystallographic waters and the Mg^2+^ cation were retained as present.

Conversely, the starting point to model the HVR in K-Ras4B and the equilibrated phospholipid membrane was a representative snapshot from one of the simulations by Prakash, Gorfe, and coworkers^34^ featuring: K-Ras4B with a full farnesylated HVR region (Figure 1a; residues 167- 185); a phospholipid membrane (95 POPS; 319 POPC); and aqueous NaCl solution. From this equilibrated structure, we deleted all ions and water molecules except those falling within 5 Å of any phospholipid or HVR atom, so as not to disrupt equilibrated conditions in the vicinity of the membrane and HVR. Subsequently, to merge the crystalline GTP-bound G domain and its crystallographic waters (*i.e.*, 6vjj)^33^ with the equilibrated HVR, we superimposed backbone heavy atoms of residues 166-167 in both species, then deleted residues up to and including His166 in Prakash’ structure, as well as residue Lys167 in 6vjj;^33^ we note that the G domain in 6vjj deviates little from the simulated one (RMSD 0.785 Å). At this stage, for simplicity, we also capped the *C-* terminal Fcy185 with an *N-*methyl group (rather than the *O-*methyl group that is present in the mature form).^37^

After superimposing and merging, the *reduce* utility in *AmberTools* (v. 19)^70^ was employed to add hydrogens to K-Ras4B G domain residues; predict histidine tautomerization (on Nε2 in all cases and with no positively charged histidines); model optimal orientations of Asn/Gln sidechains (ignoring the aforementioned catalytic Gln61); and confirm the absence of disulfide bridges. The *PropKa* package^71^ predicted all residues to be in their standard protonation states at physiological pH. Finally, the *tleap* utility^70^ was employed to model missing atoms in Arg73, and to (re)solvate K-Ras4B and the equilibrated membrane by reintroducing missing water molecules and randomly placing appropriate numbers of ions to neutralize the overall charge and restore the former 0.1 M ionic strength, with a final tally of one Mg^2+^, 230 Na^+^, and 135 Cl^-^. The resulting system (Figure S1a) retains its original dimensions^34^ (*i.e.*, a 117 × 115 × 158 Å cuboidal box), with the membrane parallel to the *xy* plane, and charged phospholipid heads about 78 and 20 Å away either box edge along the *z*-axis. The pdb file issued from *tleap*^70^ was converted to .gro^72^ format using the *pdb2gmx* utility: starting coordinates are available as Supporting Information.

### Forcefield Parameters

For reasons of mutual compatibility with other parameters, all standard K-Ras4B amino acids and terminal caps were modeled using the *ff99SB* forcefield^73^ in its *ILDN* improvement,^74^ whereas parameters for Fcy185 were based on the work by Khoury *et al*.^75^ With regard to ions, Na^+^ and Cl^-^ were treated with parameters by Joung and Cheatham,^76^ while parameters by Allnér and coworkers^77^ were used to model Mg^2+^. For GTP and GDP (the latter present in D-NEMD simulations only; *vide infra*), we adopted the forcefield reported by Meagher *et al.*^78^ Similarly, to simulate the inorganic phosphate anion [H_2_PO_4_]^-^ present in D-NEMD simulations only, we introduced parameters by Kashefolgheta and Vila-Verde.^79^ Parameters for both lipids present in the membrane (POPC and POPS) were provided by appropriate extensions of the *Slipids* forcefield^80,81^ by Jämbeck and Lyubartsev. Finally, the chosen water model was TIP3P.^82^ Where necessary, parameters were converted to *GROMACS*-compatible^72^ formats using the *acpype* code,^83^ and the correct conversion of selected parameters was verified manually. Starting topologies are provided as Supporting Information; we note that despite the switch to *ff99SB-ILDN*^73,74^ from forcefields of the Charmm family originally used by Prakash and coworkers,^34^ no major structural differences were observed in the equilibration stages of the MD simulations (*cf.* next subsection): this confirms that the change of forcefield only has a limited effect on our simulated system.

### Molecular Dynamics Simulations at Equilibrium

Equilibrium MD simulations were carried out using the *GROMACS* package (version 2021.5),^72^ in 20 independent 250 ns replicas (atomic velocities assigned with different random seeds). Each replica was preceded by a full steepest descent structural minimization for about 2000 steps (*i.e.*, until all atomic forces dropped below a 1000 kJ mol^-1^ nm^-1^ threshold to machine precision); and by a 200 ps equilibration stage in the *NVT* and *NpT* ensembles wherein restraints were imposed on certain atoms (details and conditions provided as Supporting Information). The 250 ns production stage for each replica, wherefore all restraints were lifted, was conducted in the *NpT* ensemble (*T* = 300 K; *p* = 1 bar), with a 2 fs timestep—lengthened from the preproduction stages, see Supporting Information—applied to the leap-frog integrator.^84^ The 300 K temperature was enforced by the velocity-rescaling thermostat,^85^ to which (1) protein + GTP + Mg^2+^; (2) membrane; and (3) H_2_O + Na^+^ + Cl^-^ were coupled separately with a 100 fs time constant. To enforce the 1 bar pressure during production, we applied Berendsen’s barostat^86^ with a 2 ps time constant: pressure coupling was applied semi-isotropically, with the *xy* plane containing the membrane coupled separately from the *z*-axis. A 12 Å cutoff was employed for the calculation of Lennard-Jones and Coulomb interactions, and for determining the closest neighbors around each atom (lists were updated every 10 integrator steps). The *Particle Mesh Ewald* (PME) method^87^ was used to calculate the Coulomb interactions, switching to reciprocal space beyond 12 Å. Lennard-Jones interactions were directly calculated up to 12 Å, and set to zero beyond this limit: effects of this were compensated, as *per GROMACS* implementation, by adding average corrections to the energy and pressure. All non-water bonds containing hydrogens were constrained with the *LINCS* algorithm;^88^ water bonds were constrained using *SETTLE*.^89^ All unspecified details were set to *GROMACS* defaults;^72^ all input files are provided electronically as Supporting Information.

### Distance Fluctuation Analysis (DF)

Broadly speaking, distance fluctuation (DF) analysis^14,16,60^ assesses whether, across an equilibrium MD trajectory or metatrajectory, all individual residue pairs in a simulated protein are moving in a more coordinated (more allosteric) or more uncoordinated (less allosteric) fashion. For a simulation of a protein composed of *N* residues, one thus typically obtains a single *N* × *N* **DF** matrix, whose individual elements *DF_ij_* represent the average degree of coordination (“DF score”) between the *i*^th^ and *j*^th^ residues. Each such element is given by the formula:

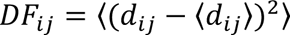

where *d_ij_* represents the distance between Cα atoms of the *i*^th^ and *j*^th^ residues in a particular MD frame, and values enclosed in ⟨⟩ denote averages over a whole trajectory or concatenated trajectories; (*i.e.*, *d_ij_* – ⟨*d_ij_*⟩ measures the deviation in each frame from the average distance observed throughout the simulation). There follows from this that residue pairs with a high DF score move in a more uncoordinated way and are less likely to be allosterically related; *vice versa*, residue pairs exhibiting low DF scores are moving concertedly and are deemed to be allosterically connected.

DF analsysis on equilibrium MD simulations of K-Ras4B in this work was conducted using our *ad hoc* code,^90^ directly on our 20 replicas (minus the first 5 ns of each), concatenated into a single metatrajectory; the procedure does not require any fitting or realignment.

### Shortest Path Map (SPM)

The theory behind SPMs (shortest path maps) is explained and justified in more detail by Osuna elsewhere:^30^ the approach is based on transforming a protein into a graph of nodes-and-edges (one node = one residue), wherein edges connecting nodes *i* and *j* (the *i*^th^ and *j*^th^ residues) are only drawn if the residues’ Cα atoms remain on average closer than 6 Å, and are given weighted lengths *l_ij_*

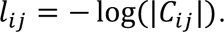

Thus, if the motion of vicinal residues *i* and *j* is more highly correlated or anticorrelated (*i.e.*, if their correlation *C_ij_*approaches +1 or –1), edges connecting them will be shorter (“heavier”), signaling a greater “transfer of information” between them.

Mathematically, correlation *C_ij_* between residues *i* and *j* across one or more MD trajectories can be computed as follows:

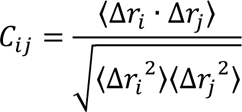

thus accounting for the average displacements of Cα atoms of residues *i* and *j* (Δ*r_i_* and Δ*r_j_*) from their positions in their protein’s most populated conformational cluster. (As for DF equations, ⟨⟩ denote averages over a whole trajectory or concatenated trajectories).

The (rather complex) node-and-edge graph is first elaborated by Osuna *et al.*’s *DynaComm.py* code,^12,30^ which then simplifies it as follows. First, the code determines the shortest possible path (*i.e.*, along the heaviest/shortest edge or succession of edges) from each residue to every other residue: to give an idea, K-Ras4B, with its 184 residues, features 16652 shortest paths. At the end of the process, edges most often traveled through in these residue-to-residue paths are given greater weights than those that are seldom or never passed, with the most often used edge acquiring a normalized weight of 1: a final SPM is then produced showing residues linked by edges whose normalized weights exceed an arbitrary 0.3 threshold.

To produce a SPM, *DynaComm.py* only requires average distance matrices and correlation matrices. Conveniently, in our case these could be directly derived from our equilibrium MD metatrajectory (*i.e.*, the 20 concatenated replicas, in .xtc format) using the “matrix” command in the *cpptraj* tool,^91^ after a few trivial realignment steps and a clustering procedure based on Cα atoms: these steps are detailed in Supporting Information. As Supporting Information, we also provide the necessary *cpptraj* input files to perform all these steps—including clustering and matrix derivation, the matrices themselves, and the output generated by *DynaComm.py*.

### Dynamical Nonequilibrium Molecular Dynamics (D-NEMD)

Short D-NEMD simulations (6 ps perturbation + 44 ps production when surviving to completion; *vide infra* and Figure 2) were conducted starting from 71706 individual frames (“windows”) directly isolated from our metatrajectory with atomic velocities (stored every 50 ps; Figure 2, thick black lines). Such frames (73.1% of the total number saved with velocities) were chosen because they were deemed to be loosely “reactive” based on the presence of a nucleophilic water in the vicinity of the γ-phosphate in GTP, and on Lys16 forming a hydrogen bond with one of the Oγ atoms of GTP (these criteria were inspired by a previous QM/MM study,^40^ and are explained in full in Supporting Information, wherein we also provide exact “reactive” frame counts per replica).

**Figure 2.**
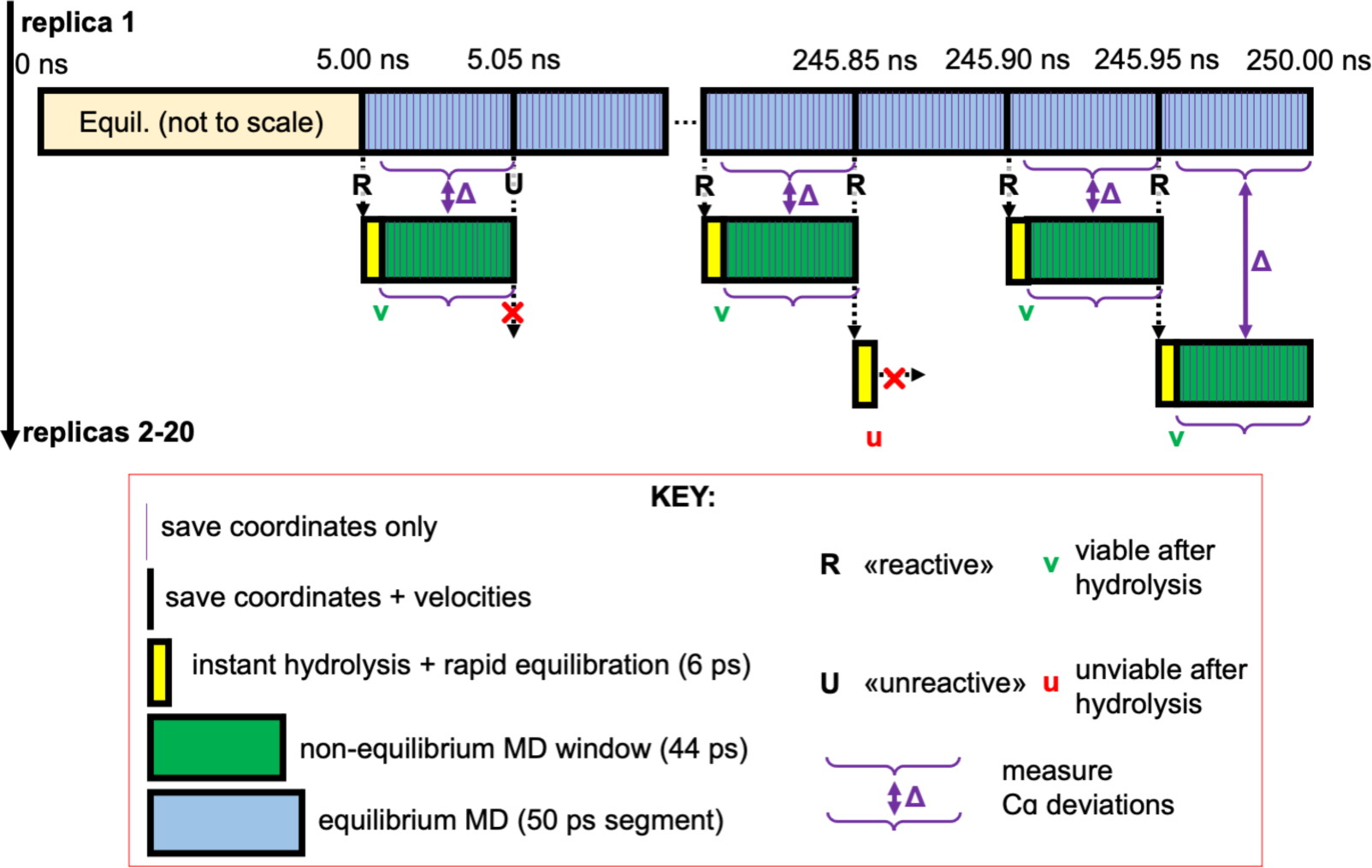
Overview of D-NEMD simulations setup, exemplified for equilibrium MD replica 1 and repeated for the remaining 19 replicas. Barring an initial 5 ns period (labeled “Equil.”; pale orange box), poses saved with velocities every 50 ps in the 5 to 245.95 ns interval (thick black lines) are assessed based on loose “reactive” (R) or “unreactive” (U) character, based on the vicinity of the nucleophilic water to GTP:Pγ (*cf.* Supporting Information). From each “R” pose, a new D-NEMD window is initiated after instant hydrolysis of GTP (see main text and Supporting Information). After perturbation (6 ps; yellow boxes), windows developing unviable product geometry (u) are discarded, while D-NEMD production (44 ps; green boxes) is initiated for viable geometries (v). Deviation of Cα atoms between equilibrium and non-equilibrium MD at equivalent points in time is then monitored across each window pair, and averaged across all windows and replicas to obtain final per-residue values at regular intervals over 50 ps.

The concept behind D-NEMD, which is extensively reviewed by Oliveira *et al.*,^17^ is illustrated in Figure 2: an identical perturbation is instantaneously introduced in each isolated window, after which MD is resumed for a short period, and the structural response of the protein is monitored over time. The structural responses at equivalent points in time are then averaged for all D-NEMD windows. In our specific case, the (chemically coherent) perturbation introduced in each window entails the immediate conversion of GTP^4–^ and nucleophilic water into GDP^3–^ and a free [H_2_PO_4_]^-^ anion at their forcefield-dictated^78,79^ equilibrium geometries. This charge-preserving interconversion is achieved by reordering and shifting the positions of as few as four atoms (details and illustration in Supporting Information, Figure S2), without altering any of the atomic velocities issuing from equilibrium MD.

Starting from this new perturbed topology (provided electronically as Supporting Information), each window is first simulated for 6 ps, with an ultra-short timestep of 0.6 fs, and *LINCS*^88^ and *SETTLE*^89^ constraints temporarily lifted (all else remains equal to the production stage of equilibrium MD; input file provided electronically). Continuing only if [H_2_PO_4_]^-^ has retained its correct geometry after equilibration (the verification process is detailed in Supporting Information),^92^ we then restore the same conditions present in equilibrium MD and continue simulating each window for a further 44 ps, reaching 50 ps (Figure 2; green boxes; input file provided electronically). This ensures that deviation in Cα atoms with respect to equilibrium can be monitored in each window every 2 ps, from 6 to 50 ps. Note that no superimpositions or realignments are required prior to measuring this deviation.

Both the D-NEMD production and perturbation stages—the latter indicated as “slow growth” in previous literature,^62,63^ where it was applied by gradually switching parameters between an ATP and ADP+inorganic phosphate states—come at a small loss, with as many as 80.6% of the initial “reactive” windows surviving to completion. This loss, which is due unphysical breakup of the reconstructed [H_2_PO_4_]^-^ in a minority of windows because of instantaneously shifting charges, still leaves us with a statistically sound combined D-NEMD production time of over 2.54 µs (44 ps × 57823 windows). Exact totals are provided in Table S1. Statistical validity was confirmed by quantifying standard errors of the mean for each of the 184 Cα atom deviations, at 50 ps after hydrolysis, which revealed very small errors ranging from ±0.0028 to ±0.0053 Å (data not shown).

### Anisotropic Thermal Diffusion (ATD)

ATD MD simulations were conducted starting from a set of 4900 frames isolated from our 20 replicas at regular 1 ns intervals, excluding the first 5 ns of each replica. Each frame is equilibrated (supercooled) for 200 ns at 10 K, first in the *NVT* then in the *NpT* ensemble (input provided as Supporting Information), with the cutoff for Lennard-Jones (and Coulomb interactions in direct space *per* the PME method)^87^ retained at 12 Å (δ*t* = 1 fs); the neighbor list was updated every 10 integrator steps. As for equilibrium MD simulations, pressure coupling was applied semi-isotropically, with the *xy* plane containing the membrane coupled separately from the *z*-axis, at 1 bar, again using the Parrinello-Rahman barostat with a time constant of 2 ps (as in the *NpT* equilibration phase). The 10 K temperature was retained with the velocity-rescaling thermostat,^71^ to which protein, membrane, and solute/solvent were coupled separately with a 100 fs time constant.

After equilibration at 10 K, we initiated 4900 production runs in the same *NpT* conditions at 10 K, except for GTP, which was instantaneously heated to 300 K (velocity-rescaling thermostat, time constant switched to 200 fs; input provided as Supporting Information),^71^ and the barostat, which was switched to Berendsen’s^86^ as *per* the equilibrium MD production phase. The protein, remaining at 10 K, was entirely decoupled from the thermostat. Anisotropic thermal diffusion is then derived by computing per-residue RMSD of backbone heavy atoms, after structure-fitting (RMSD alignment) on backbone heavy atoms of the first frame in each ATD production run (ignoring HVR residues).

## Results and Discussion

We will here begin by separately presenting and discussing results from the four chosen allostery detection approaches, with only cursory references to any important similarities and differences between methods and their findings as we go along. A more systematic evaluation of consistencies and differences is provided thereafter, together with contextualization and comparison to existing computational and experimental understanding of K-Ras4B. There finally follows a brief critical discussion on the key implications of our findings.

### Distance Fluctuation Analysis on Equilibrium MD

We previously showed and validated experimentally^93^ that *higher* DF scores denote residue pairs moving *less* concertedly and thus with lower allosteric dialogue; conversely, *lower* scores indicate residue pairs exhibiting *greater* allosteric coordination. We begin by analyzing the DF score matrix (Figure 3; bottom; with secondary structure elements marked along its axes), roughly in order of increasing allosteric relevance.

**Figure 3.**
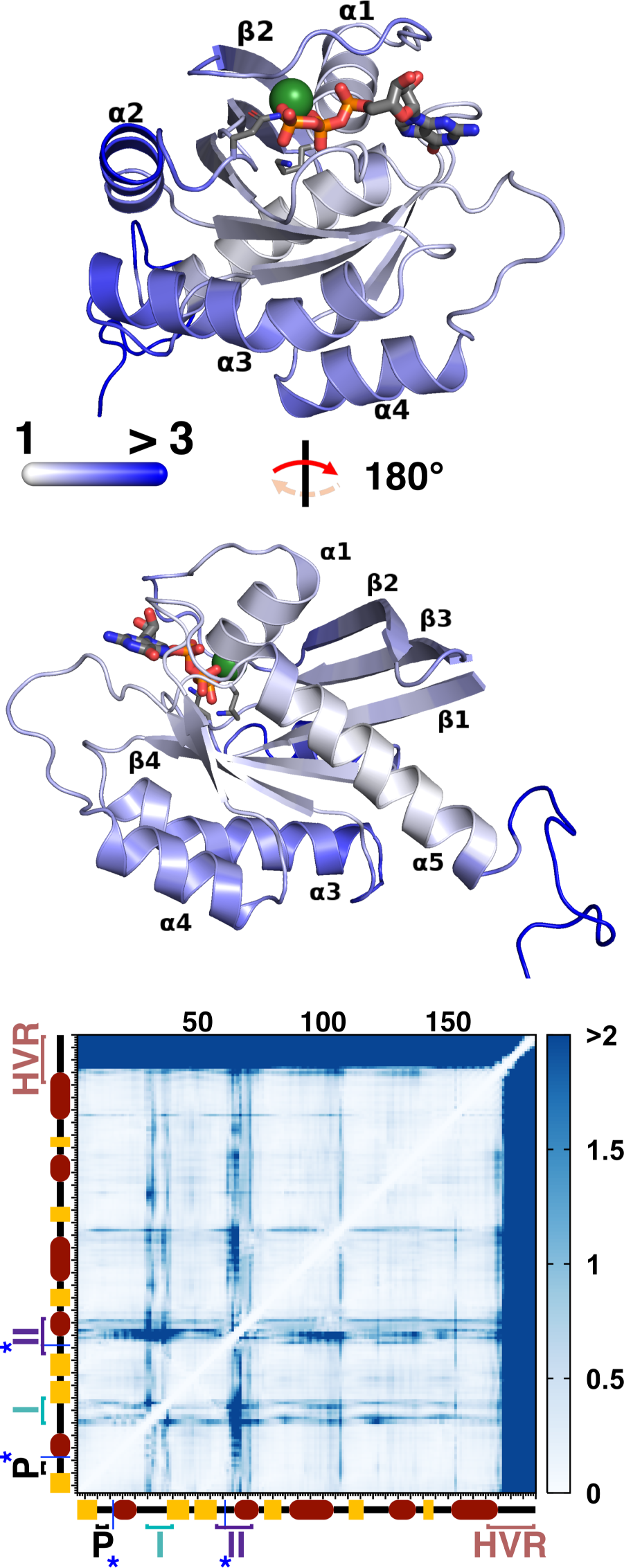
(top) Projection onto the starting K-Ras4B structure (Figure 1a) of the average (1D) DF score “felt” by each residue (see text for details), with backbone colored in increasingly darker shades of blue for increasingly higher DF scores. (middle) The same structure, rotated 180° about the axis parallel to the plane of the page. In both views: GTP is rendered as ticker sticks; K16 and Q61 as thinner sticks; Mg^2+^ as a sphere; while H, solvent, Cl^-^, Na^+^, and membrane are omitted for clarity; color code for atoms is identical to Figure 1a. Where possible, secondary structure is labeled explicitly. (bottom) 2D matrix of pairwise DF scores. Note the different scale and tones of blue. Secondary structure elements and salient conformational regions of K-Ras4B (P-loop; switches I and II; HVR) are marked along both axes for reference, as are catalytic residues K16 and Q61 (blue asterisks/lines).

Analysis shows a well-defined trend whereby the unstructured HVR (residues 169-185) is moving in a generally uncoordinated fashion with respect to the G domain (*cf.* intense blue stripes in the top and right parts of the matrix). Other salient regions with significantly low allosteric coordination to the rest of the G domain (bluer tones) include most of switch I and the portion of switch II just downstream of catalytic^40^ Gln61, including the part comprised in helix α2.

Interestingly, Thr35 on switch I and residues Thr58-Gln61 on switch II are notable outliers, with comparatively greater allosteric coupling to the G domain (whiter stripes). These outliers are all essential for reactivity: Thr58, Ala59 and Gln61 are involved in hydrolysis regulation,^40^ whereas Thr35 completes the coordination shell of the GTP-chelating Mg^2+^ (and loses coordination once hydrolysis deactivates switch I).^44^ Higher allosteric coupling to the bulk of the G domain is also observed for Lys16 (another catalytically relevant residue)^40^ and for the mutation-prone P-loop.

Finally, areas with the highest allosteric relevance in the DF matrix are spanned by regions of secondary structure, with the exception of helix α2 and the *N-*terminus of sheet β2 (due to their partial locations on switches II and I, respectively). In particular, central β-sheets β4 to β6 and helix α5 span the “whitest” areas in the matrix, suggesting they may form a coordinated and compact hub, with a high degree of inter-residue crosstalk that is crucial for the repartition of allosteric signals to other K-Ras4B regions.

To aid in the interpretation of the DF matrix in Figure 3, as Supporting Information we provide a video that dynamically maps its information onto the structure of K-Ras4B: in a stepwise progression from residue 2 to residue 185, the video highlights the degree of coordination of each residue with every other residue, with projected colors on the backbone evolving accordingly. In addition to this video, in the top part of Figure 3, we have projected a 1D-averaged version of the DF matrix onto our starting K-Ras4B structure (*cf.* Figure 1a), which is also represented just beneath it (Figure 3; middle) after an 180° rotation. This 1D-version of the matrix is obtained by summing DF scores in each matrix column (*i.e.*, residue by residue), and averaging the resulting score over the 184 pairs formable by each residue including with itself. In doing so, one gains insight into the “average” degree of allosteric (un)coordination experienced by each residue even if the two-dimensionality of the matrix is lost. Besides clearly recapitulating all of the allosteric traits emerging from the 2D matrix, flattening the matrix in such a way, for example, further brings to light the fact that helices α3 and α4 and sheets β1 to β3 are less allosterically coordinated on average (bluer) than helix α5 and sheets β4 to β6 forming the central allosteric hub (Figure 3; middle structure). In addition, this operation provides per-residue DF scores that make for an easy comparison with the other allostery detection methods.

To summarize, 1D and 2D DF data alike concur in finding sheets β4–β6 to be particularly coordinated in their movements, and thus exhibiting high allosteric intercommunication; the same relevance is observed for helix α5. Most of the G domain only retains a moderate degree of internal coordination, with the P-loop; possibly sheets β1, β3 and most of β2; catalytic Lys16 as part of helix α1; and (to a lesser extent) Gln61 and very minor portions of switches I and II falling in this category. In contrast, the HVR and the greater part of effector switches I and II are found to have a very low degree of allosteric coupling with other regions, exhibiting clearly uncoordinated movements.

### SPM derived from equilibrium MD

The shortest path map (SPM) representing the main allosteric communication route^12,30^ across K-Ras4B, as derived from our full metatrajectory, is illustrated in Figure 4 as purple spheres (residues) connected by black lines (paths). The *PyMOL* session file used to derive Figure 4 (left) is also provided along with all SPM output and input, as Supporting Information. As a reminder, the (dimensionless) “shortness” of each path segment is proportional to its thickness in Figure 4, and proportional to the intensity of allosteric communication between the two vicinal residues it connects.

**Figure 4.**
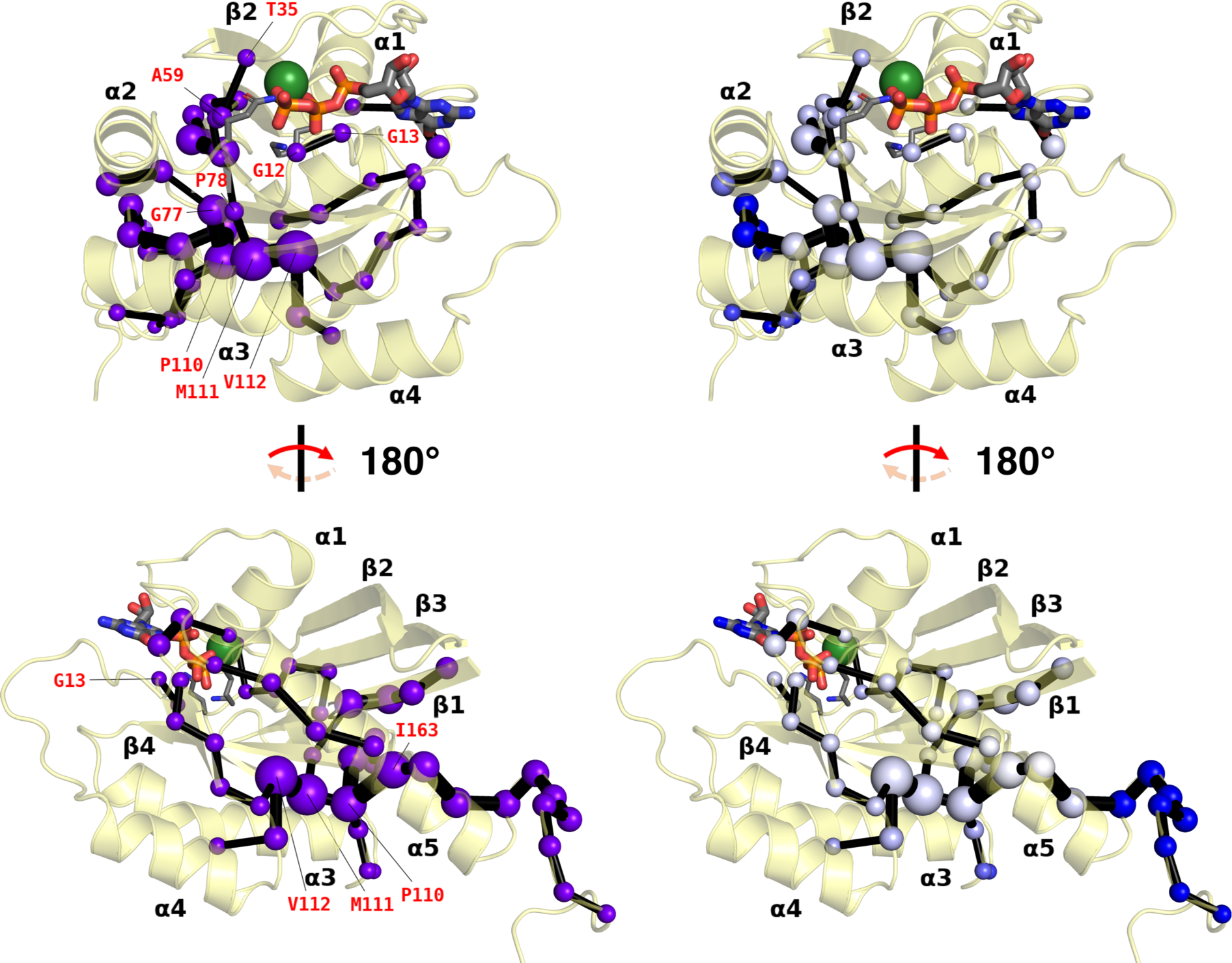
Allosteric shortest path map (SPM) calculated from our 20 concatenated equilibrium MD simulations of K-Ras4B. On the left, we show the simple SPM drawn on the starting structure: purple spheres represent communicating residues (Cα atoms) and thickness of black sticks is proportional to their “allosteric closeness” (see text); where possible, residues mentioned in the main text are labeled in red. On the right, we color the SPM according to the 1D DF score of residues that are present (*cf.* Figure 3, top panels). Views and atom color codes are identical to those for Figure 3, with added transparency.

The SPM paints a very similar picture to DF analysis, as commented later in this subsection. Indeed, the SPM features a prominent allosteric pathway that originates midway along the HVR (Figure 4; bottom)—close to the membrane anchor point and best describable in the *C-* to *N-* direction—that eventually branches out to reach most allosterically relevant G domain regions, while virtually avoiding (most of) switches I and II. Working its way up the HVR and the *C-* terminal half of the α5 helix, the path first reaches Ile163; Ile163 communicates very strongly with residues Val112, Met111 at the *N-*terminus of sheet β5, and Pro110 on the α3–β5 loop, with an elevated relative shortness of 0.79 out of 1 (Figure 4; bottom; *cf. Computational Methods*). Indeed, by virtue of this shortness, Ile163 and the three α3–β5/β5 residues are seen to form the main allosteric communication hub in K-Ras4B: in particular, the path connecting Val112 and Met111 themselves is the shortest one in the protein.

Five shortest path branches depart from this four-residue hub. We label these I to V and discuss them in detail in the Supporting Information. From an allosteric point of view, Branch V is the most intensely travelled route and the one spanning the most interesting regions. It branches off to β4 from the hub on β5, with a dual coupling to from Pro110 (β5) to Gly77 (β4) and from Met111 to Phe78 (β4); from there it further branches out into three subbranches V.1, V.2, and V.3 (*cf.* Supporting Information), of which V.3 is the most allosterically interesting, since it reaches all crucial areas of the active site. More specifically, moving through Val7 on β1 and the *C-*terminal residues of β3, it reaches all the way up to residues Thr58 and Ala59 on switch II, which as mentioned during DF analysis are amongst residues controlling hydrolysis.^40^ Equally intriguingly, and again in line with DF analysis, V.3 also encompasses Thr35, which keeps Mg^2+^ coordination; as well as the oft-mutating Gly12 and Gly13 in the P-loop. In fact, the importance of some of the most crucial residues along the SPM is also confirmed by mutagenesis studies:^41^ we comment on this in more depth later on in this section.

In short, the shortest path originates from the HVR: this likely picks up allosteric messages from the membrane and conveys them to a major hub located on helix α5, loop α3–β5, and the *N-* terminus of sheet β5. From this hub, the path branches out to fully encompass sheets β1, β4, and β6. Crucially, key areas within the binding site or its vicinity are also eventually reached, most notably Thr58 and Ala59 on switch II; Thr35 on switch I; and the P-loop. On the other hand, regions such as helix α3, the remainder of switches I and II (minus Thr58 and Ala59), and the remainder of sheet β2, remain out of reach.

Indeed, consistency of SPM findings with DF data is readily ascertainable (Figure 4; right), particularly in terms of the 1D average DF scores projected on the structure in the top and middle of Figure 3. With the sole exception of the HVR, which only the SPM identifies as allosterically relevant, all areas touched by the SPM are also found to be allosterically important by our DF analysis, both in terms of mutual allosteric crosstalk and in terms of (1D) average coordination (Figure 3). This is clearly demonstrated by the fact that blue spheres on the right of Figure 4 (*i.e.* high average DF scores, poor allosteric coupling) are clearly limited to HVR residues. *Vice versa,* all remaining areas with low average coordination coincide with those untouched by the SPM.

### D-NEMD simulations

D-NEMD simulations do not assess allostery under equilibrium, unlike SPM or DF analysis, but they instead model its role in relaying information away from the nucleotide binding site upon forced GTP hydrolysis to GDP and [H_2_PO_4_]^-^. When mapped onto the initial K-Ras4B structure, for example 36 ps after hydrolysis (Figure 5), the average structural response of the protein reveals a very interesting hydrolysis propagation pattern. We should note that the pattern’s progression is uniform throughout the monitored 50 ps hydrolysis period: to follow it in full from 0 to 50 ps, the reader is referred to the video we provide as Supporting Information.

**Figure 5.**
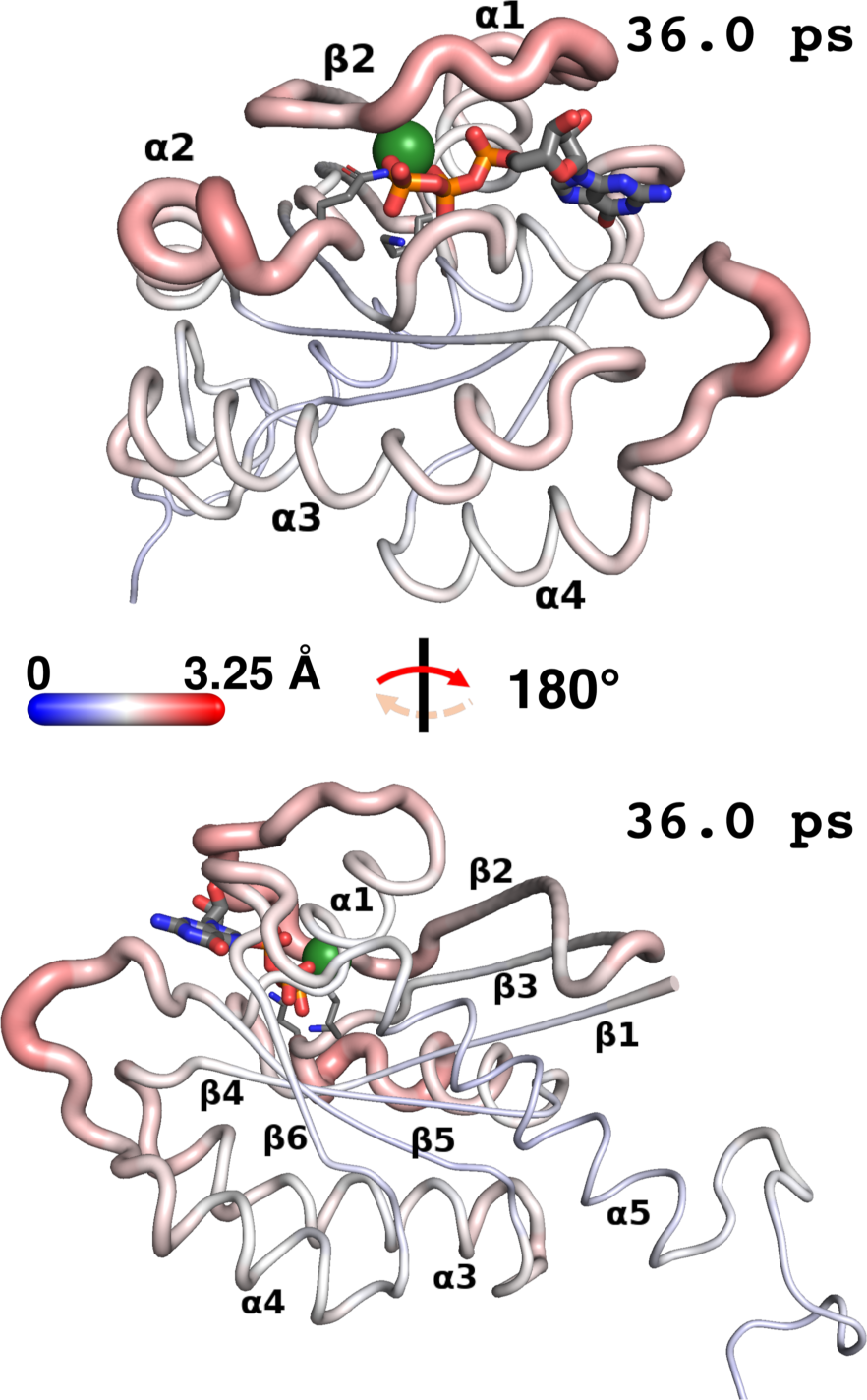
D-NEMD simulation results viewed on the starting K-Ras4B structure (views and atom color codes are identical to those for Figure 3). Backbone thickness of each residue and intensity of red *vs.* blue is proportional to the average direct deviation (in Å) of its Cα atom in D-NEMD simulations—*i.e.*, after forced GTP hydrolysis—from its position in equilibrium MD simulations at an equivalent moment in time. Deviations are averaged over 57823 “reactive frames” surviving hydrolysis (Supporting Information; Table S1); in this Figure they are exemplified as they appear 36.0 ps after hydrolysis, but they were measured up to 50.0 ps after (see video in Supporting Information). The choice of time is the one that visually maximizes contrast between most- and least-deviating residues at the chosen color scale (0 to 3.25 Å deviation); note however, that no residue reaches either of these values at the chosen time.

In any case, areas in the immediate proximity of the binding site evidently feel the greatest effects from hydrolysis (redder color = stronger deviation in Figure 5), but not in a uniform way: it is clear (Figure 5; top) that switches I and II in their entirety bear a greater brunt. In fact, with a 2.27 Å deviation at 36 ps, Glu63 on switch II is the residue with the greatest average deviation compared to equilibrium MD, and catalytic Gln61 itself deviates by 2.01 Å. By comparison, less deviation is observed in other areas adjacent to the GTP binding site, namely the β6–α5 loop and the P-loop, whose average deviations of 1.79 Å and 1.74 Å, respectively, make them appear much whiter in Figure 5 (top; the β6–α5 loop appears behind the guanine moiety of GTP). Catalytic residue Lys16, despite its proximity to the cleaving phosphate, is basically unaffected in the first 50 ps after hydrolysis.

Compared to equilibrium MD, sheet β1, core sheets β4–β6, and helix α5 (Figure 5; bottom) exhibit an even lower combined average deviation at 36 ps of 1.48 Å: this echoes their rigidity in SPM and DF findings, and suggests that while they could act as rigid transit hubs for the transmission of allosteric signals, they are themselves unaffected by hydrolysis. Focusing on remaining peripheral regions in K-Ras4B, it is equally clear from Figure 5 that not all of them are affected by D-NEMD hydrolysis in the same way: as in previous D-NEMD studies,^62-65^ greater flexibility and/or solvent exposure is by no means synonymous with a greater deviation upon perturbation and, indeed, eliminating most deviational “noise” that is not directly linked to the introduced perturbation is precisely one of the specific advantages of D-NEMD. The only distal region from the hydrolyzing phosphate to undergo significant deviation, though still part of the binding site, is evidently the β5–α4 loop (on the right of Figure 5; top; and on the left of Figure 5; bottom), on a par with deviations observed for residues in switch II; remaining peripheral regions do experience some of the effects of hydrolysis after 36 ps (*cf.* paler red areas in Figure 5), but not to the same magnitude; these include, *e.g.*, the α1 helix (≤1.98 Å) in its farthest part from GTP, the β2–β3 loop (≤1.95 Å), the *N-* and *C-*termini of helix α3 (≤1.89 Å), the *N-*terminus of helix α4 (≤1.87 Å), and the β2 sheet outside switch I (≤1.79 Å). Deemed allosterically important by the SPM approach but recognized by DF analysis as one of the K-Ras4B regions with the greatest flexibility and solvent exposure, the (very peripheral) HVR stands out for its particularly low deviation (≤1.64 Å), entirely comparable to the α5/β1/β4–β6 core: this again suggests it is likely unaffected by allosteric signals emanating from the active site.

The reader will recall (*cf. Computational Methods*) that D-NEMD statistics were collected on 57823 loosely “reactive” windows surviving hydrolysis. In defining these, we purposely ignored the conformation of Gln61, which in reality is crucial to properly lock the nucleophile into position for attack,^40^ since it would have lowered the number of viable frames. Still, we can confirm that if, for completeness, perturbations are recalculated on this fraction of surviving D-NEMD frames in which Gln61 too is suitably positioned for hydrolysis (23209; criteria in Supporting Information), the result is virtually indistinguishable from Figure 5 (data not shown but available upon request). To recapitulate, D-NEMD data show that the effects of the forced hydrolysis of GTP have immediate and significant repercussions on switches I and II. Other areas in the vicinity of the nucleotide either feel these effects considerably less, including the P-loop and the β6–α5 loop, while catalytic Lys16 is barely affected. Away from the active site, the rigid α5/β1/β4–β6 core and the HVR are largely unaffected by hydrolysis; whereas some of its effects do indeed propagate to other peripheral regions, most prominently to the β5–α4 loop.

### ATD simulations

Like D-NEMD simulations, anisotropic thermal diffusion (ATD) simulations too assess propagation of allosteric crosstalk patterns upon perturbing our system from equilibrium. In this case, however, the perturbation does not consist in forced GTP hydrolysis, but on reheating GTP alone to 300 K after supercooling the entire system at 10 K, as repeated starting from 4900 individual frames isolated from the MD metatrajectory. Allosteric signals irradiating from the binding site in ATD simulations are therefore conceptually different compared to those in D-NEMD simulations: they should capture areas of K-Ras4B that are allosterically dialoguing with GTP as a whole rather than those sensing the effects of GTP hydrolysis. Also, unlike in D-NEMD, progression of allostery in ATD is measured with respect to the first production frame in each window (whence all atoms will have shifted by some degree), *not* with respect to an exactly equivalent time at equilibrium (when shifts are only concentrated in perturbed areas): this means that noise resulting from ordinary K-Ras4B flexibility is not entirely canceled. Quite on the contrary, we found that including the HVR when aligning to the first frame of an individual window always led to generally high levels of “RMSD noise” on all the K-Ras4B structure, making it hard to extract meaningful allosteric information. For this reason, we proceeded to exclude most of the HVR (residue 169 and above) from alignment to the first frame in each ATD window and exclude it from the ensuing ATD analysis (Figure 6).

**Figure 6.**
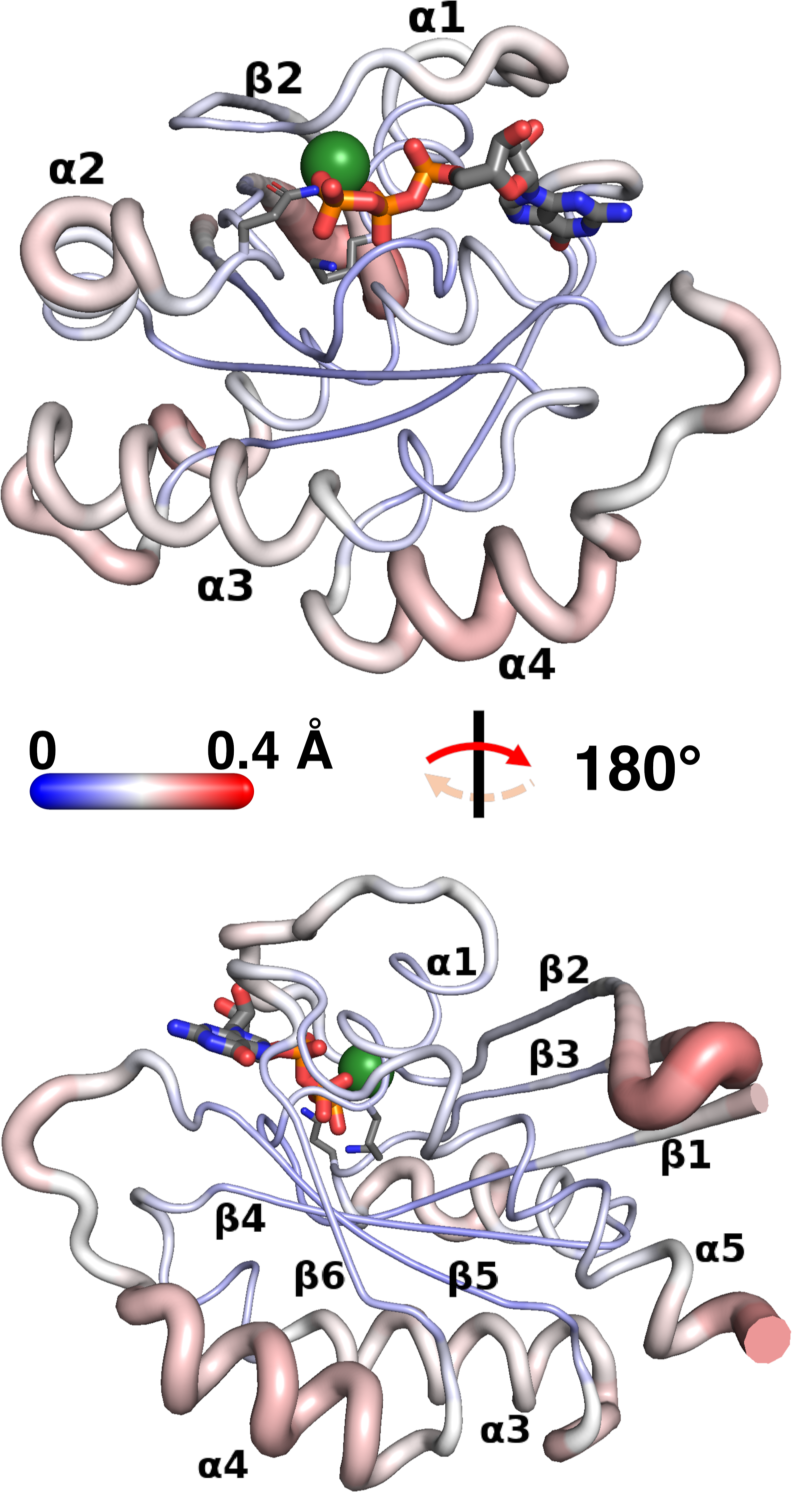
ATD results viewed on the starting K-Ras4B structure (views and atom color codes are identical to those for Figure 3; note, however, that in this case analysis was truncated beyond residue 168, and therefore most of the HVR is omitted). Backbone thickness of each residue and intensity of red *vs.* blue is proportional to the average RMSD (in Å) reached by that residue by the end of an ATD production run compared to its beginning. Averages are taken over the 4900 individual frames on which we performed ATD simulations.

ATD simulation data projected on the starting K-Ras4B structure (Figure 6) show that while areas of greater average RMSD (redder) with respect to the start of GTP heating generally coincide with areas of greater absolute deviation in D-NEMD simulations after hydrolysis (Figure 5), extents by which they deviate in the two situations can be quite distinct. The most patent difference is the more moderate deviation in switches I and II compared to other areas: this is both in contrast to D-NEMD simulations, whereby they are the part of K-Ras4B that is most affected by hydrolysis (Figure 5), and in contrast to their uncoordinated (flexible) status detected in the SPM (Figure 4) and by DF analysis (Figure 3). More concretely, the maximum average RMSD detected upon reheating GTP is 0.22 Å (Asp30) in switch I and 0.23 Å (Ala66) in switch II / helix α2 (Figure 6; top): even visually, it is clear that there are areas in K-Ras4B that *per* Figure 6 show greater or comparable average deviation, all of which are far from the GRP binding site. The first of these is β2–β3 loop (Figure 6; bottom), whose Gly48 exhibits the greatest deviation of all residues considered (0.29 Å), followed by helix α4, with an average deviation of 0.24 Å. The α3–β5 loop, whose residue 107 incidentally stands out as allosterically uncoupled in the DF analysis (Figure 3), shows a comparable degree of average deviation (Figure 6; bottom), and the β5–α4 loop deviates just a little less, at 0.18 Å on average (Figure 6; top and bottom).

The only other region of the active site that shows appreciable deviation—which, incidentally, is found by D-NEMD too—is the β6–α5 loop, also at 0.18 Å (Figure 6; top; behind the guanine moiety in GTP). Conversely, what stands out in the active site is the absence of significant deviation in the P-loop (Figure 6; top), and in catalytic residues Gln61 (Figure 6; top) and Lys16 (Figure 6; bottom), suggesting that while coupled to GTP once it is hydrolyzing (D-NEMD), these areas are not relevantly coupled to GTP prior to it.

Focusing, finally, on the least-deviating parts of K-Ras4B away from the active site (Figure 6), we once again confirm that the α5/β1/β4–β6 core is minimally perturbed, excluding the final few *C-*terminal residues of α5, which begin to feel the strong deviation experienced by the rest of the (unincluded) HVR group.

In summary, areas that are perturbed upon heating GTP in ATD simulations qualitatively overlap with areas feeling the effects of GTP hydrolysis in D-NEMD simulations. However, it is interesting to note that all perturbed areas distal to the binding site experience a greater average deviation compared to those in its proximity, including switches I and II. The P-loop and catalytic residues Gln61 and Lys16 are significant outliers, showing no allosteric coupling to GTP heating at all; the central α5/β1/β4–β6 core is also found to be unaffected by GTP heating, but in this case much expectedly.

### Listening to the Various Languages

While clear trends already emerge from isolated analyses in Figure 3–Figure 6, in order to obtain meaningful information about K-Ras4B it is of course necessary to compare the four allostery detection methods more systematically: this is possible through the heatmap plotted in Figure 7. In the heatmap, which only spans residues 2-168 since ATD was not meaningfully analyzable in the HVR, we compare a distinctive raw per-residue score *S_raw_* chosen for each allosteric language, rescaled/normalized so that the score *S_norm,i_* for the *i*^th^ residue will always fall between 0 (black) and 1 (yellow).

**Figure 7.**
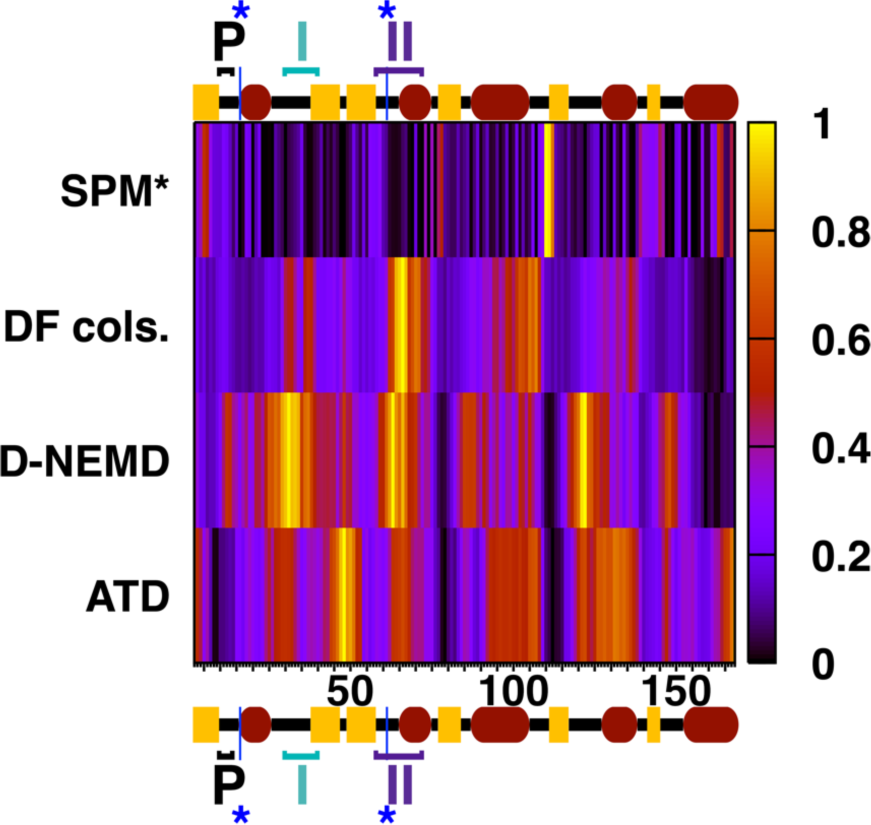
A combined overview of the four allosteric languages compared in this work, with per-residue representative scores for each method (see main text for chosen ones) normalized to between 0 (black) and 1 (yellow), passing through shades of purple, red, and orange. Since the ATD analysis excludes the HVR, all comparisons have been truncated to span residues 2-168 only: secondary structure elements, the P-loop, and switches I and II have been marked along the top and bottom axes, with the two blue (*) denoting positions of catalytically relevant Lys16 and Gln61. *The chosen SPM score (see main text) denotes greater residue rigidity when closer to 1, and is presented in the top row to distinguish it from other method scores, which indicate more rigidity when closer to 0.

Full details about derivation of *S_norm_* scores from *S_raw_* are provided as Supporting Information. Regarding the process of choosing a suitable per-residue *S_raw_* for each method, ATD and D-NEMD (Figure 7; two bottom rows) already provide per-residue scores (respectively, RMS backbone deviation and Cα deviation from equilibrium). *S_raw_* for DF analysis (Figure 7; second row) are simply taken to be average per-residue DF scores emerging from the flattening of the 2D matrix (*i.e.*, those projected on the structure in Figure 3). For SPM (top row of Figure 7), we choose a *S_raw_*that is derivable from the *DynaComm.py* code (SPM prominence), and is proportional to the number of times that a given residue is ‘visited’ as an intermediate point along one of the many individual *shortest paths* that connect each residue to every other residue.

Of note, per-residue *S_norm_* scores for SPM are conceptually opposite to the other three chosen scores. In the former case, residues with the highest *S_norm_* correspond by definition to those falling along the final SPM (Figure 4): residues with scores tending to 1 represent therefore those with the highest degree of allosteric communication. Conversely, in the other three languages, scores tending to 1 either denote residues with *low* allosteric communication/rigidity or those with the greatest deviation upon heating or hydrolysis. The top row of Figure 7 therefore follows an opposite chromatic trend compared to the bottom three.

### Interpreting the Various Languages and Speaking the Languages of Experiment

With this *caveat* in mind, Figure 7 unequivocally indicates that there is a generally high degree of consensus between all four approaches in categorizing several important regions of K-Ras4B. At the same time, consensus is clearly not universal, and there are some differences between certain languages: these differences are, incidentally, entirely expectable, since they directly reflect the fact that, as set out in the *Introduction*, not all allosteric routes are created equal.^6^ More specifically, there will be certain pathways that a system will preferentially be prone to explore only when at equilibrium, by virtue of its intrinsic dynamics; there will be other pathways whose exploration will only be able to gather some pace after a biochemical trigger (in our case, GTP hydrolysis); and there will be allosteric pathways that remain significant in both circumstances, possibly to different extents. It is precisely due to the different declensions of allostery that we here invoke the use of more than one language in the first place. A full portrait of the complex allostery of K-Ras4B in which the role of each residue is meticulously reconstructed is only possible, as we shall see, thanks to the unique nuances that each allostery detection method is able to provide, interrogating K-Ras4B on its propensity to visit allosteric states that become relevant at very different stages of its biochemical lifecycle.

Broadly speaking, the SPM and the DF analysis are natural partners (akin to linguistic cognates) requiring no additional information other than the original set of unbiased MD simulations from which they are constructed. In our specific case, they inform us from slightly different perspectives on likely allosteric hotspots characterizing the GTP-active state. Additionally, by virtue of their two-dimensional nature (Figure 3; bottom) DF scores also allow one to break down general allosteric signals into components of “individual” allosteric dialogue between specific residues or groups of residues. Even if our system does not undergo major conformational rearrangements during MD, the SPM and DF analysis uncover allosteric hotspots that are likely to mediate such rearrangements at sufficiently long timescales, and those that would be most disruptive if interfered with.

ATD and D-NEMD also revolve around the same set of unbiased MD simulations but, each in its own way, they inform us on allostery by monitoring the average *change* in dynamics upon introduction of a specific perturbative event. They identify “temporary hotspots” that, at a given point in time, will harbor the greatest repercussions form the allosteric perturbation in question. Allosteric propagation routes (responsive residues) can in principle be indirectly deduced by retracing the perturbation backwards or forwards in time, and thus distinguished from regions whose dynamics are unaffected simply because they are not involved in conveying that particular perturbation. By artificially heating GTP in a supercooled K-Ras4B, the more ‘unphysical’ ATD reveals allosteric signals specifically emanating from the active site. D-NEMD similarly informs explicitly on temporary allosteric hotspots at (a) given time(s) after perturbation, but is based on instantaneous differences resulting from a real biochemical event (phosphate cleavage) with tangible biochemical consequences (specifically, which areas are most likely to intercept signals from hydrolysis and impair the effector-recruiting interfaces characterizing the GTP-active state). In the subsections that follow, we dissect trends in the normalized scores shown in Figure 7, relating them, entirely *a posteriori*, to the most recent experimental and computational allosteric understanding of K-Ras4B in its active state.^37,41,44,66^ This represents the most significant final validation emerging from the integration of our methods. Where possible, we will stress which areas of the protein show universal consensus across languages, and which ones do not and why.

### Predicting an Allosterically Compact G domain

The most obvious consensus across all normalized scores in Figure 7 is that the highest allosteric coupling at equilibrium (*i.e.*, SPM prominence; light) and either some very low fluctuations (*i.e.*, DF; dark) or low susceptibility to perturbation (*i.e.*, D-NEMD, ATD; dark) are invariably observed at the cusp of the α3–β5 loop and β5 sheet or, otherwise put, at the hub centered on residues 110-112 whose emergence from the data we have frequently referred to in preceding subsections. Such a clear-cut consensus strongly proposes these residues as the main allosteric core in K-Ras4B. A similar consensus is observed at the *N-*terminus of the β4 sheet, reconfirming that this sheet too is allosterically crucial, including for the propagation of allosteric perturbations.

Helix α5 also emerges as a strong contributor to internal allostery at equilibrium that is generally unscathed by GTP hydrolysis, except at its *N-*terminus. Three approaches confirm it as a fundamental allosteric player: average DF scores are the lowest (blackest) in the entire protein, signaling strong allosteric coupling; SPM shows high ‘closeness’ values (red bands) for crucial residues Ile163, Arg164, and Lys167; whereas D-NEMD shows moderate perturbations at similar positions along the helix. Only ATD scores (lighter than expected in Figure 7) clearly disagree in this case, likely due to contamination from HVR noise.

Finally, the β6 sheet, also part of the central core, is universally recognized as an allosterically ‘intermediate’ area (Figure 7 scores in the ‘purple’ range, due to uneven *S_norm_* distribution). It should be envisaged as both retaining some degree of allosteric relevance at equilibrium but— while not nearly comparable to the most perturbed areas—it is not as immune to allosteric impulses from GTP hydrolysis or heating as other parts of the G domain that feel no effect.

In short, our ‘linguistic evidence’ describes vital elements of the G domain as forming an entity that is fairly compact allosterically: consistency of this picture with available experimental data is evident.^41,44^ Going further in depth, the conspicuous study by Weng *et al.* mentioned in the introduction,^41^ thanks to point mutations systematically introduced across the entire K-Ras4B structure, specifically speaks of anisotropic allosteric coupling across the central beta sheets.

More encouragingly still, when the authors^41^ go on to dissect which positions are most allosterically affected by mutations, they provide an explicit list of 8 ‘novel’ allosteric nodes at various locations in the G domain (on top of 10 in the vicinity of the binding pocket): strikingly, while not all of these residues are comprised in the regions of universal consensus mentioned so far from Figure 7, 6 out of 8 of these allosteric residues are actually comprised in the SPM, on which we mark them (Figure 8) in purple and ruby red. Experimentally identified residues lying on the SPM are Val7 (β1), Thr58, Ala59 (switch II), and—most importantly—Pro110 (α3–β5), Phe141 (β6), and Ile163 (α5), which precisely emerge in the SPM as some of the “closest” (most communicating) residues within our consensually identified allosteric hub. In addition, a seventh site, Asp54 on β3, while absent from the SPM, interacts electrostatically with SPM node Lys5 (β1) (pink in Figure 8); Gly10 at the *N-*terminus of the P-loop is the only relevant allosteric site that is neither part of the SPM nor in contact with one of its residues. A further experimental confirmation of the allosteric importance of helix α5, besides the appearance of Ile163 in Figure 8, comes from studies of pan-mutant inhibitor BI-2865^56^ (*cf. Introduction*), wherein certain evidently ‘allostery-reactivating’ mutations in helix α5 are seen to diminish its effects despite the inhibitor—which favors the inactive GDP state—binding far from the helix.

**Figure 8.**
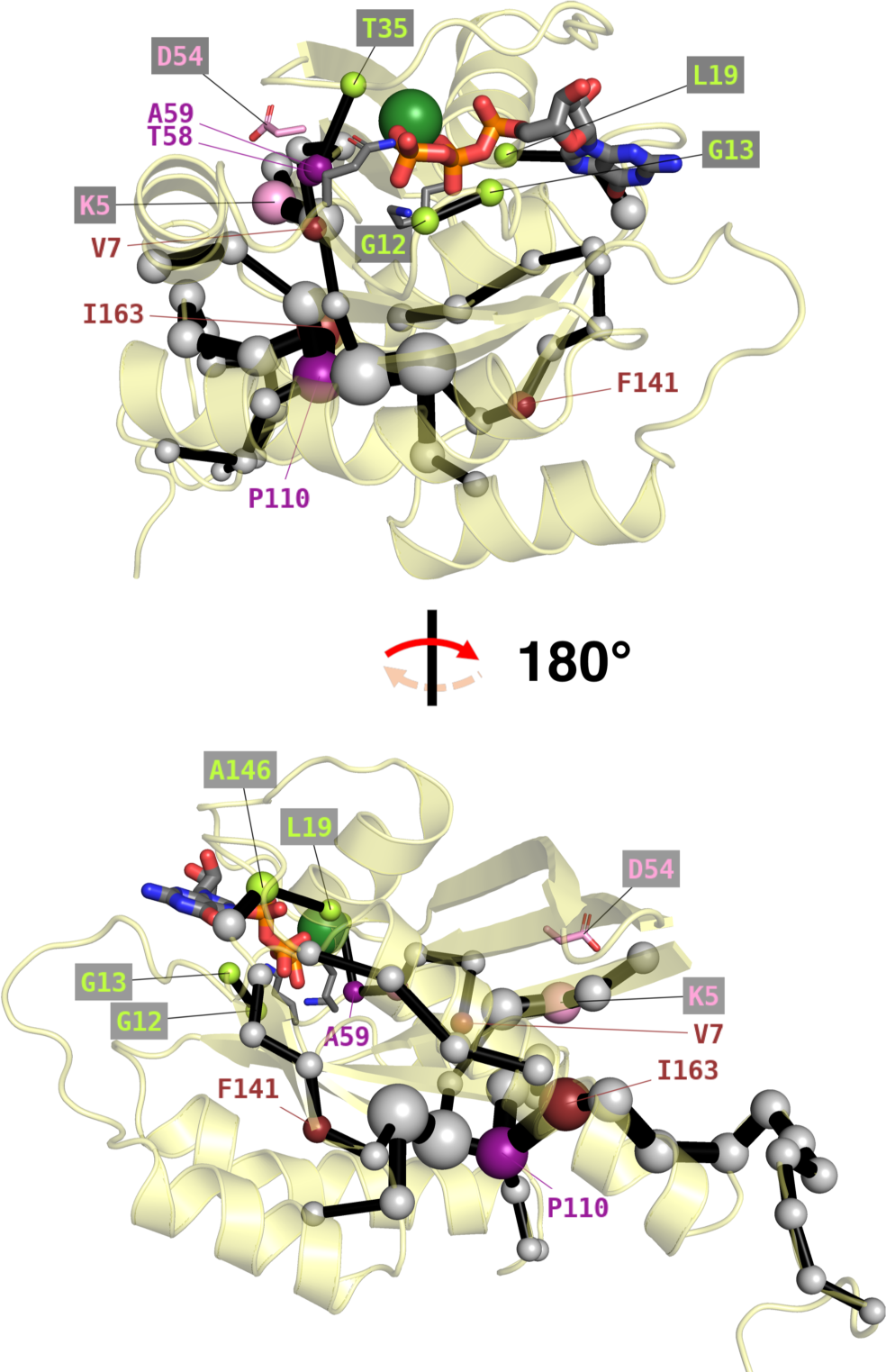
Alternative view of the SPM (Figure 4) from identical angles, but with 7 out of 8 ‘novel’ allosteric residues uncovered experimentally (the 8^th^, Gly10, is omitted).^41^ The 6 that directly appear on the SPM are either labeled in purple, if they also correspond to known cancerous mutations,^42^ or in ruby red otherwise. Asp54, which does not lie on the SPM but interacts with Lys5 (pink), is rendered as sticks and itself labeled in pink. In lime green, we label other SPM residues closer to the active site which frequently mutate in cancers.^42^ All other SPM residues are denoted by off-white spheres.

For completeness, we should assess how the other languages identify sheet β1, which contains the experimentally and SPM-relevant residues Val7 and Lys5, we should note that *per* Figure 7, remaining allostery detection methods (*i.e.*, ATD, D-NEMD, and DF analysis) paint a picture that is less clear-cut than that for the α3–β5 loop, helix α5, or sheets β5 and β6. Allosteric coupling detected by DF is not as strong as for other elements of the allosteric hub and definitely in contrast with the prominence predicted by the SPM; similarly, perturbation ‘felt’ upon heating GTP is certainly more accentuated than expected. Only the D-NEMD method confirms that there is not as much perturbation after GTP hydrolysis.

Consensus of our non-SPM methods for regions containing the remaining residues highlighted in Figure 8 is discussed over the next two subsections.

### Predicting Cancerous Mutations on the P-loop and Outside the Switches

In fact, if we turn our focus onto the most abundant^42^ clinically relevant mutations of K-Ras4B, we find that the recolored SPM in Figure 8 holds yet more interesting aspects to reveal. The first thing to note is that three of the ‘novel allosteric nodes’ described by Weng *et al.*^41^ actually also correspond to frequently documented cancerous mutations: these are the ones marked in purple in Figure 8, and are Thr58, Ala59, and Pro110. (Specifically, the former two suffer most often point mutations to Ile58 or Thr59, while the latter frequently suffers a complex frameshift mutation).^42^

More interestingly still, beyond these ‘novel allosteric residues’ the SPM also captures several of the other clinically documented mutation sites, albeit with prominences that are not as pronounced; these are all marked in lime green in Figure 8. The first of these sites to note are, of course, Gly12 and Gly13 on the P-loop, whose cancerous mutations especially to Asp12, Asp13, and Cys13 are by far the most abundant:^37,42^ they feature in one of the SPM branches, but with a modest prominence that approaches the 0.3/1 ‘shortness’ threshold below which residues are excluded. With a similarly ‘low shortness’, we also show in lime green three additional clinically relevant mutation sites featured on the SPM (occurring at a much lower frequency^42^ than those on the P-loop): Leu19 on helix α1, responsible for binding the guanosine moiety of GTP; Thr35 on switch I, responsible for maintaining Mg^2+^ coordination; and Ala146 on the β6–α5 loop, which is part of a G5 motif that is conserved across the RAS family^39,56^ (E*x*SAK, residues 143-147).

For the sake of our *a posteriori* comparison, we should thus reextend our discussion to assess how the remaining three allostery detection methods narrate the story of these residues and their regions (*i.e.*, back to the *S*_norm_ heatmap in Figure 7). (Switch I and Thr35 are discussed in the next subsection).

For the P-loop, DF and D-NEMD match the SPM in painting a profile of ‘moderate allosteric relevance’ that is similar to the one for the β6 sheet discussed previously. At equilibrium (DF matrix; Figure 3), there is clearly some degree of allosteric communication with the G domain, as signaled by low (white) scores: this is also clear from the DF video in Supporting Information (residues 10-14). What is perhaps striking in this case is the much lower-than-expected deviation upon GTP heating signaled by *S_norm_* scores for ATD (black band in Figure 7, bottom row; blue in Figure 6): this signals that prior to hydrolysis, the P-loop is only really allosterically communicating with the rest of the G domain but not with GTP, despite its proximity. Only upon hydrolysis, D-NEMD scores in Figure 7 suggest that there is some coupling between the P-loop and the cleaving P_i_, though not on the same level as the switches.

Similarly, for the Leu19 region of helix α1 and the Ala 146 region of the β6–α5 loop (G5 motif), we have a situation of relatively high allosteric communication at equilibrium *per* the DF scores (Figure 3; Figure 7) that matches moderate SPM prominence and, in this case, both ATD and D-NEMD agree that there is a fair degree of perturbation upon GTP heating or hydrolysis.

In reflecting on these findings, we should recall that the role of cancerous mutations is to enhance the GTP-active state in some way: this is quite the opposite of obliterating its allosteric communication pathways like the other mutations in Figure 8 (complex mutations on Pro110 are a more complicated case). In this respect, the nuances painted by our four allostery detection methods for Gly12, Gly13, Leu19, and Ala146 are encouragingly in line with this picture: their moderate allosteric relevance at equilibrium in a G domain that is much more dependent on the β- sheet/α5 allosteric hub for its integrity ensures that they can mutate without disrupting the GTP-active state. At the same time, their prominent coupling to GTP and their perturbation upon hydrolysis suggests that mutation of these residues—particularly Leu19 and Ala146 that are on the guanosine end of the active site—could enhance activity by mitigating the allosteric perturbations that characterize K-Ras4B inactivation. In fact, similarly to the ‘reactivating mutation’ of helix α5 that hampered BI-2865 activity by inactivating GTP hydrolysis,^56^ the G5 motif is also observed to undergo such mutations.^56^

In the case of P-loop, which is close to the triphosphate moiety of GTP and not as connected to it prior to hydrolysis, one could also envisage a role as one of the allosteric hubs that controls reactivity: it would normally receive instructions from the G domain to initiate GTP hydrolysis, but P-loop mutations could disrupt these instructions and impair reactivity. In the case of mutations to Asp12 and Asp13, effects on hydrolysis could even be purely electronic in nature, with a much more ‘direct’ influence on reactivity.

In general, direct electronic or electrostatic intervention as opposed to subtle allosteric disruption of the active site could well be the most likely mode of action of other key carcinogenic mutations on switches I and II. We will more articulately speculate on these possibilities later, when we focus on allosteric control of catalytic residues.

### Predicting the Allosteric Role of Switches I and II

Having mentioned mutations on switches I and II, which as we know represent the most obvious trait of K-Ras4B distinguishing the GTP-active from the GDP-inactive state (Figure 1), it is now a good point to focus our discussion on how our simulations identify these crucial regions. In this case too, we find that all four methods provide very eloquent findings: indeed, the majority of switches I and II show very low allosteric coupling to the G domain core at equilibrium (Figure 7; dark SPM; light DF), and are clearly identified by ATD and D-NEMD alike as allosteric hotspots for perturbative events (lightest shades, bottom two rows). Significantly, however, there is one subsection on each switch that ostensibly bucks these trends, showing up as distinctly colored bands in Figure 7 for all four languages: on switch I, this is the central portion that comprises the clinically^42^ (and catalytically)^40^ crucial Thr35 (Figure 8), whereas on switch II, it is the *N-* terminus that harbors the clinically^42^ (and catalytically)^40^ crucial Thr58, Ala59 (Figure 8) and Gln61. These residues retain a modest degree of allosteric coupling to the allosteric hub: all have comparatively lower DF scores, and all except Gln61 feature in the SPM.

Consistency of these findings with experimental evidence is patent. The substantial flexibility of most of switches I and II even at equilibrium is entirely in line with the reported existence of several conformational states even in the GTP-bound state^36,37,39,44,53^ that ultimately enable K- Ras4B to recruit regulatory proteins such as RAF1 and GAP. Still, while switches I and II remain (expectedly) very mobile in the GTP-active state, they both retain mild allosteric coupling to the G domain in catalytically crucial regions (*cf.* Figure 3; bottom): Mg^2+^-bound Thr35 in the former case, and Thr58/Ala59/[Gln61] in the latter case. On top of being the defining trait that allows switches to form functional interfaces only in the active state, it also confers them control on GTP hydrolysis itself. At the same time, both switches are clearly predicted to be very perturbed by hydrolysis: this unequivocally suggests that once hydrolysis does occur, and departure of P_i_ and Mg^2+^ severs all allosteric links, switches look set to lose their ability to attain active-state conformations (eventually reaching the status in Figure 1c), and their affinity for effectors will be compromised. In other words, our simulations confirm that there is a clear connection between GTP hydrolysis and switch deactivation without the need of reproducing a major conformational change in either switch.

### Allosteric Control on GTP Hydrolysis

We have seen that catalytically relevant residues Thr35, Thr58, Ala59, and Gln61, which are also subject to pathogenic mutations,^42^ are located on special subportions of switches I and II that all our chosen languages depict with fairly similar characteristics. Similarly, we have already discussed our consistent findings for mutation-prone Gly12 and Gly13, which are located in the P-loop in the vicinity of the triphosphate moiety of GTP. To these residues, we should add the final catalytic actor, Lys16, which is just downstream of the P-loop, and which we have explicitly marked from Figure 3 to Figure 7. A good reference for the hydrolytic mechanism, including on the role played by GAP and its Arg789 (Figure S1b),^69^ is the QM/MM investigation by Calixto *et al.*^40^ Prior to hydrolysis, Gln61 helps position the nucleophilic water to attack GTP:Pγ, and both prior to and during hydrolysis, cationic Lys16 lowers the free energy barrier by abstracting negative charge from the transition state. One would expect Lys16 be detected to be at least as allosterically important at equilibrium as Gln61, if not as Thr58 and Ala59, in the context of the active site preorganizing for hydrolysis; yet, Lys16 remains excluded from the SPM, with a *S_norm_* that is even less prominent than that of Gln61. The only allosterically relevant aspect of Lys16 is general allosteric coupling with the G domain detected in the DF matrix (Figure 3; bottom, Supplementary Video). This suggests that while crucial for catalysis, Lys16 did not evolve to become as prominent an allosteric regulator of it.^40^

With Gly12, Gly13, Thr35, Thr58 and Ala59 featuring in the SPM (Figure 8) but at the same time Lys16 and Gln61 excluded from it, and with DF scores showing a moderate degree of allosteric coupling to GTP at equilibrium, it is difficult to univocally establish exactly how each mutation will hamper GTP hydrolysis, or indeed, in unmutated K-Ras4B, which of these residues will mostly control hydrolysis through electronic/electrostatic intervention as opposed to acting through subtle allosteric regulation of the active site.

Of course, it is not implausible that known^42^ mutations of Gly12, Gly13, or Gln61 to Asp would be more likely to intervene electronically, as in all three cases the Asp would be within reach of sequestering Arg789 in GAP (Figure S1b)^69^ from GTP:Pγ. Similarly, mutation of Thr35 to Ala^42^ could impede Mg^2+^ binding, thus leaving GTP unchelated and unable to hydrolyze. As also recognized by the QM/MM study,^40^ Gly13 and Ala59 too can exert an electrostatic influence on the reaction barrier and therefore on K-Ras4B reactivity. On the other hand, vicinal residues such as Gly12 and Thr58, both of which feature on the SPM but do not have ascertained electrostatic roles, could indeed be controlling reactivity through allosteric effects on the active site alone. Proving these hypotheses, however, would require additional simulations and experiments and is beyond the scope of this work.

### Predicting Allosteric Control of Other Functional Interfaces

Since switches I and II are the main effector recruiters in GTP-active K-Ras4B, it is of course clearly encouraging to see that D-NEMD simulations (Figure 5, Figure 7) predict both switches to be the two areas in K-Ras4B that are most perturbed by GTP hydrolysis; this is because these perturbations are synonymous with incipient interface degradation.

In fact, we observe that the agreement between D-NEMD simulations and key structurally known interfaces formed by K-Ras4B is much more far-reaching. *E.g.*, taking the interface with effector RAF1 as an example (PDB ID: 6xi7; Figure S1c),^33^ we see that Asp153 on the *N-*terminus of helix α5 forms an integral part of that interface; in full agreement with this, the reader will recall that the *N-*terminus of the otherwise unperturbed helix α5 (*vide supra*) is the only part of the helix for which D-NEMD simulations predict some degree of perturbations. Further still, in all remaining areas of that interface outside switch I—namely, helix α1, the remainder of sheet β2, loops α1–β2 and β2–β3—D-NEMD simulations (Figure 5, Figure 7) also predict large perturbations *post-*hydrolysis, again showing that the event impairs all parts of the interface and is consistent with obliterated RAF1 binding by the GDP-inactive state.

While we cannot review all known K-Ras4B–effector interfaces in full, we can see that, in fact, in five other interfaces dissected by Weng *et al.*,^41^ the six residues that are in common to all and are most impacted by their mutation (Gln25, Asp33, Ile36, Asp38-Tyr40; marked in Figure S1c) are located in helix α1, loop α1–β2, and sheet β2, signaling that the perturbation of this wider area during hydrolysis (and not just switch I) has actual biological repercussions; indeed, an inactive switch I (Figure 1c) also entails open β2 and β3 sheets.^37^ The ability of D-NEMD to capture interface perturbations in general^17^ is also reconfirmed through these simulations.

Even the interface with GAP (Figure S1b; PDB ID: 1wq1),^69^ which is not an effector but oversees hydrolysis and therefore evidently requires preference for the GTP-active state, spans this perturbed α1/α1–β2/β2 area that includes switch I. In addition (and this of course is in common with other effectors), GAP also interfaces (Figure 5, Figure 7, Figure S1b) with the equally perturbed switch II and loop β3–α2, and other more marginally perturbed but crucial areas including, notably, the P-loop (wherein mutations could therefore have a sterically disruptive effect for this deactivating interaction on top of the direct electronic effects discussed earlier).

### Predicting the α3–β5 Loop as a Secondary Allosteric Switch

We have amply seen that Pro110 (α3–β5), Met111 (β5), Val112 (β5), and Ile163 (α5) are of cardinal allosteric importance for maintaining the structure of the G domain (Figure 4); unsurprisingly, importance of loop α3–β5 and helix α5 is highlighted in a number of experimental contexts,^33,39,41^ a number of which— particularly for helix α5—we have already discussed.

On top of harboring allosteric hub residue Pro110 at its *C-*terminus, the *N-*terminus of the α3– β5 loop is a secondary allosteric switch that regulates the ‘kink’ in helix α3 (*cf.*, *e.g.*, Figure 3), which in turns helps keep switch II in the active state alongside GTP.^39^ In this respect, it is very interesting to notice that in Figure 7, the *N-* and *C-*terminal portions of the α3–β5 loop are recognized with very different characters. Obviously, as part of the allosteric hub, the *C-*terminal portion of the α3–β5 loop resembles the β5 sheet itself; on the other hand, the immediately preceding *N-*terminal portion of the loop, along with helix α3 itself, exhibits radically opposite characteristics (Figure 7), *i.e.*, low allosteric communication at equilibrium (lighter DF, darker SPM), but clearly more sensitivity to hydrolysis (lighter D-NEMD). These very distinct behaviors are entirely compatible with the reported^39^ allosteric roles of helix α3 and loop α3–β5 as a secondary allosteric switch: the *C-*terminal portion of α3–β5 keeps switch II in place in the active state; however, once the *N-*terminal portion of the loop and helix α3 itself feel the effects of GTP hydrolysis, this ‘gatekeeper’ effect is lost.

### The HVR—Flexible but Fundamental

Ile163 on helix α5 is the residue that first intercepts incoming allosteric signals from the HVR. Speaking of the HVR itself, while its presence in Figure 7 is limited to its first few residues because of the noise associated with ATD (which we will therefore ignore), representations in Figure 3–Figure 5 show that it too is approached quite differently by our chosen languages. While DF analysis points to high flexibility / poor allosteric coordination (intense blue color in Figure 3), SPM predictions go in the opposite direction (Figure 4; Figure 8) suggesting HVR can play an important allosteric role despite its flexibility. After hydrolysis, in principle, over the 50 ps they cover in this work, D-NEMD simulations detect very little deviation in the HVR: on top of being a testament to the better ability of D-NEMD to cancel out non-allosteric noise, this intriguingly suggests that allosteric perturbations from hydrolysis are not wasted by dissipating back into the membrane.

All of the above findings are nicely consistent with experiment: if, on the one hand, flexibility is in line with irresolution of the HVR in most of the available crystal structures,^37^ allosteric dialogue with the G domain is equally in agreement with experimental reports.^36^ In terms of reported^45,46^ HVR-switch (and G-domain—membrane) contacts, we should note that these are not replicated in our simulations, wherein centers of mass of both components remain distant in all our replicas, in the 35-40 Å range. Nonetheless, throughout our MD simulations we do observe a high degree of K-Ras4B mobility across the surface of the lipid bilayer: this fits in with the established role^36^ of K-Ras4B in recruiting effector monomers to the cellular membrane and bringing them together so that they can dimerize.

### Predicting Allosteric Pockets

We should finally extend our analysis to the four allosteric pockets I to IV previously recognized by Grant *et al.* through simulations^43^ and later validated experimentally,^41^ in order to evaluate how well our own recipe is able to recognize these pockets and thus assess its potential in predicting allosteric pockets in new systems. (In the designation used by both studies,^41,43^ inhibitors Sotorasib and BI-2865 both bind to pocket II). While of course interference with all four pockets ultimately ushers in diminished K-Ras4B activity, our own assessment reveals that pocket II,^43^ pocket IV,^43^ and pockets I/III^43^ fall, respectively, into three distinct categories of which the latter is conceptually distinct from the former two.

More specifically, pocket II should be envisaged as a stabilizer of the GDP-inactive state (Figure 1c), which was indeed considered alongside the GTP state in Grant *et al.*’s simulations;^43^ the pocket occupies the volume created between inactivated switch II and allosteric helix α3. The shallow pocket IV is found^43^ through blind docking on GTP-active K-Ras4B, and occupies the volume between switch I, helix α1, and the rest of sheet β2. The reader will recall that these regions correspond to the interface with RAF1 discussed earlier (Figure S1c) and indeed, the purpose of targeting this pocket would be to disrupt the K-Ras4B–RAF1 interface. In this case, since disruption of the interface is amply predicted by D-NEMD, one could assume that application of our approaches to other systems could automatically help identify other interfaces that could be therapeutically targeted. On the contrary, however, we must point out that, strictly for the purposes of our discussion, identification in a system of pockets such as pocket IV would require some prior degree of knowledge about interfaces in the first place, to understand which interfaces are therapeutically targetable and which are not. Similarly, identifying pockets like pocket II would require structural knowledge of other biologically relevant conformational states, and explicitly taking them into account.

On the other hand, pockets such as I/III specifically disrupt structurally crucial allosteric sites within the GTP-active state. This means that they are fully identifiable by our chosen approach and simulations: as clearly inferable especially from SPM (Figure 4) and D-NEMD data (Figure 5), both these pockets occupy regions of the utmost allosteric importance at equilibrium in the active state, while feeling little to no perturbation from GTP hydrolysis. More specifically, pocket I is located in the coupled region between sheets β1, β3, and the *C-*terminus of helix α5; and pocket III spans none other than the main α3–β5/β5/α5 allosteric hub. Interference with these pockets is thus bound to bring maximum disruption to the active state, and experimental data certainly suggests so too.^41^ This correspondence with experiment bodes well for novel systems whose allosteric properties are not as pervasively known: in such cases, one could for example compare and contrast the information provided by the SPM and D-NEMD to predict binding sites that could disrupt allostery if interfered with; in systems that are allosterically less compact, and wherein areas of allosteric coupling and decoupling are more easily distinguishable, DF analysis has certainly proven to be a useful guide too.^60^

### The Final Picture: The Membrane Talks to GTP, GTP Talks to the Switches

In summary, after piecing together information from our own simulations and from previous work, there emerges a chronologically clear, articulate, and experimentally consistent allosteric portrait of K- Ras4B. For the chosen GTP-active state of K-Ras4B^33^ (*vide infra*), languages consistently describe a compact protein in which allosteric signals appear to be far-reaching but rigorously compartmentalized: prior to hydrolysis, at equilibrium, a set of pathways affords a more or less strict control on hydrolysis itself while retaining a mild dialogue with the switches; once hydrolysis begins, on the other hand, signals mainly (but not exclusively) propagate to effector switches I and II, disrupting their function.

More specifically, at equilibrium, helix α5 and sheets β1, β4–β6, as a rigid core, constitute the allosteric centerpiece of the G domain; in particular, the high allosteric significance of sheets β4 and β5 is captured by all four allosteric languages, and that of helix α5 by three of them out of four. Despite its intrinsic flexibility, it is also coherently found that the HVR is paramount in relaying signals between the membrane and the above rigid core. While the intricate network of allosteric dialogue ostensibly avoids the majority (but crucially not the entirety) of switches I and II, there are convincing signals from SPM, and to a certain extent DF and ATD, that there does exist a direct allosteric coupling going from the HVR all the way up to key binding site residues, including catalytic ones. Said binding site residues are first and foremost the P-loop and guanine-binding residue Ala146 on the β6–α5 loop (G5 motif).

Albeit more blandly, however, parts of both switches are also involved in the allosteric control: switch II through Thr58/Ala59 (DF and SPM) and switch I through Thr35 (DF and SPM). Existence of these allosteric links at equilibrium to both switches, however mild, is the fundamental reason why switches can mediate interaction with regulatory proteins only in the active state. Similarly, the electronic relevance^40^ of some allosterically important active site residues not within switches suggests that allosteric control on GTP hydrolysis is likely to be exerted for the greater part through these residues rather than through Lys16 and Gln61, which influence the reaction more ‘directly’, through electronic effects.

Moving on along our timeline of events, anyhow, once hydrolysis is initiated, the strong allosteric link between the cleaving phosphate center and switches I and II (with Gln61)—which get visibly perturbed (DNEMD; Figure 5) and of course mediate K-Ras4B inactivation—could not be clearer. There also is an unequivocal disruption of the secondary switch on the *N-*terminus of the α3–β5 loop; of other regions forming interfaces, including the *C-*terminus of helix α1, the β2 sheet outside switch I, and loops α1–β2 and β2–β3; and, again, of the β6–α5 loop.

While in proximity to the hydrolytic center, P-loop and Lys16 remain comparatively less perturbed by hydrolysis. It is also important to recall that the central β-sheet core, α5 helix, and HVR are entirely decoupled from hydrolysis, even despite the latter’s flexibility, meaning signals from hydrolysis are unlikely to be relayed back to the membrane: this serendipitous finding actually makes biological sense, insofar as the GTPase should have evolved in such a way that little or none of the energy resulting from hydrolysis is (wastefully) dissipated away from biologically functional areas.

### The Importance of Learning many (Allosteric) Languages to Decipher Unbiased MD

One final point to consider are the reasons and implications of our methodological choices. With so many different *in silico* “allosteric languages” available across the scientific community, some now for many years (*cf. Introduction*), it was clearly beyond the scope of this work to exhaustively compare-and-contrast as many methods as possible. Rather, we have opted to limit ourselves to four methods—quite different in scope and theoretical origins—and not ‘compare’ them, but use them as extensively as possible on a well-documented oncotarget, with the aim of highlighting just how many subtleties there are to allostery, and how powerful a suitably planned combination of *in silico* allostery detection methods can be in capturing them adequately.

We should stress that our choice of methods does not imply an indication of preference or support for any one of them: in fact, we have mainly aimed to seek methods that were as heterogeneous as possible and beyond our immediate areas of expertise. Much like learning new languages costs time but is worth the investment, we recommend to authors wishing to undertake a new allosteric study that they too seek to maximize heterogeneity in their chosen methods, while monitoring key aspects such as noise, portability, computational cost, and user-friendliness.

As Supporting Information, we review some of the considerations specifically applying to our own choice of methods, and review them critically. We should obviously stress that plenty of other options would have been available for our investigations. While it is not possible to review them here in detail, they should by all means be considered as alternatives by any reader wishing to plan their own investigation. A few examples taken from across the realms of equilibrium and perturbative allostery alike include Normal Mode Analysis,^94^ Gaussian Network Models,^95^ Leverage Coupling,^21^ Perturbed Ensemble Analysis,^96^ and Networks of Local Correlated Motions.^97^ Additional methods and online resources are reviewed in more detail elsewhere.^5,21,23^ It is also important to point out that findings from a chosen combination of languages could be further reinforced by forms of systematic coevolution analysis^2,26^ and/or analysis of the most frequent pathogenic mutation sites.^42^

Returning to our own choice, and especially in light of the experimental validation made earlier, we can make a solid argument that when applied and interpreted rationally, the four allostery detection methods featured in this study—all of which are automatable and fairly portable to other systems—can work synergistically to provide a high level of allosteric detail about a particular system that they would be unable to provide separately. A further fundamental advantage of our computational recipe that is worth reiterating is its total applicability to unbiased MD simulations, without the need to sample large-scale conformational changes or start from alternative conformations: after all, the large amount of data originating from this work is, essentially, derived from just a single starting structure. Alternative approaches to study different systems should be chosen so that they retain the same advantages.

## Summary and Conclusions

Allostery has evolved alongside proteins to regulate most aspects of their biological function, enabling residues that give rise to functional interfaces, pockets and/or active sites to dynamically influence each other even when they lie tens of Ångströms apart. Perturbations of the delicate allosteric equilibria governing proteins can have far-reaching and oftentimes detrimental effects, for example ushering in pathogenic alterations of reactivity and promotion of harmful interactions over beneficial ones. The multifaceted ways in which allostery can manifest itself during the lifecycle of a protein resemble spoken “languages”: speaking them correctly (*i.e.*, modeling them accurately) should be of great help in understanding molecular mechanisms driving (changes in) biological function of a particular protein. Unbiased molecular dynamics (MD) simulations are an option, but require decryption with a suitable allostery detection method (language) due to the sometimes too long operational timescales of allostery; while several such (very valid) methods exist, they tend to be used in isolation by their reference community, thus affording a partial, “monolinguistic” picture.

In this work, we have conducted a series of 20 independent unbiased MD simulations of the fully solvated, membrane-embedded, unmutated GTPase K-Ras4B, in an “active” GTP-bound state that is characterized by two allosteric switches poised to recruit effector proteins. Aiming to prove our argument that the combination of more than one allosteric language should appreciably improve the allosteric model of a system under study, we proceed to decrypt allostery in these simulations using four representative allostery detection methods (languages). The intensely studied oncoprotein K-Ras4B,^36,37,41^ with its notoriously difficult pharmaceutical targetability, was deliberately chosen to prove the benefits of this combination of methods.

Languages chosen to interrogate the active state of K-Ras4B at equilibrium include: (i) distance fluctuation (DF) analysis,^14,16,60^ which postulates that pairs of residues moving more concertedly than others—with distances remaining closer to the simulation average—should represent hotspots of allosteric change; and (ii) the shortest path map (SPM),^12,30^ which reconstructs the main allosteric communication pathway based on networks of vicinal residues that move with high (anti-)correlation. Out of equilibrium, (iii) dynamical non-equilibrium MD (D-NEMD) simulations^17,61-66^ which spawns a large number of short (50 ps) MD simulations from as many unperturbed MD frames, in which GTP is nearly instantaneously hydrolyzed by brute force: deviation form equilibrium MD, averaged over all short simulations, reflects the degree of allosteric perturbation induced by hydrolysis. Finally, (iv) anisotropic thermal diffusion^67^ supercools a series of frames isolated at regular intervals, equilibrates them, and then reheats the GTP only: deviation with respect to the first frame reflects the degree of coupling to the active site.

As expected, the four methods paint a very articulate allosteric picture of GTP-active K-Ras4B. Owing to their ability to capture different expressions of allostery, while all four languages concur in predicting high or low allosteric importance for certain areas of K-Ras4B, they provide intriguingly different answers for certain other areas: it is thanks to these linguistic nuances that one can provide a clear chronological dimension to K-Ras4B allostery. At equilibrium, prominent allosteric communication pathways travel from the membrane and through the flexible hypervariable region, from whence they reach a paramount allosteric hub centered around the *C-* terminal half of (tendentially rigid) helix α5, sheet β5, and around loop α3–β5. From here, they branch out to encompass most remaining parts of the protein, notably including—albeit to moderate degrees—residues on several sides of the GTP binding site. Amongst these are residues on both switches (Thr35 on I and Thr58/Ala59 on II) and mutation-prone Gly12 and Gly13 on the P-loop. As shown by QM/MM reactivity studies,^40^ these residues or others in their immediate vicinity, such as Lys16 and Gln61, exert some form of control on GTP hydrolysis, signifying that the active state can withhold its own inactivation regardless of the presence of the GTPase stimulator GAP. Despite the above allosteric links, switches exhibit low allosteric coordination, in agreement with their interfacial plasticity. Upon hydrolysis, most of K-Ras4B remains unaffected, except for the two switches (on which it is known to have an inactivating impact), and the α1/α1– β2/β2 area: all belong to crystallographically known interfaces.^33,69^

Consistency with experiment is recognizable in a number of other aspects, notably in the importance of the α5/β5/α3–β5 allosteric hub,^41^ general allosteric compactness,^41,44^ and retention of allosteric links in the hypervariable region and in both switches.^37^ Our data also agree with experiment in terms of the allosteric relevance of helix α5 and the G5 domain;^56^ and two of four pockets suggested by mutagenesis^41^ which, rather than stabilizing the GDP-inactive state, are likely to disrupt the GTP-active state instead, are also located in allosterically active regions.

To conclude, while notably different in conception, pitfalls, and genesis (exactly like spoken languages), the four chosen allosteric languages in synergy have depicted a very nuanced and experimentally consistent allosteric portrait of K-Ras4B that they would have been unable to provide if applied on their own. Crucially, decryption of such an articulate portrait was possible even if our simulations were all begun from a single structure in its GTP-active state. The coherent results produced by our chosen techniques on allosteric pathways in K-Ras4B provide elegant reconfirmation of allostery as a universal property of proteins, from therapeutic targets to biocatalysts, regardless of the computational “languages” used to decipher it. We believe our work proves the benefits of applying as many “allosteric languages” as computational resources permit.

## ASSOCIATED CONTENT

### Supporting Information

The following files are available free of charge.

Depiction of the starting structure and the K-Ras4B—GAP complex; MD preproduction details; Alignment, clustering, and matrix derivation for SPM; Full SPM branch details; Further D-NEMD details and “reactive” pose counts; Derivation of *S_norm_*scores; Critical assessment of the chosen methods. (PDF)

Equilibrium MD: starting coordinates, topology, input; ATD: input; D-NEMD: topology and input; SPM: alignment scripts, input matrices, output. (ZIP)

D-NEMD: average Cα deviation from 0 to 50 ps, projected onto the starting structure. (MP4)

DF analysis: intensity of DF scores for all possible K-Ras4B residue pairs, highlighted on the starting structure, progressing from residue 2 to residue 185. (MP4)

### AUTHOR INFORMATION

## Author Contributions

SAS and GC conceived the work and planned the experiments. SAS prepared the initial topology as received from previous simulations, supervised the work, drew Figures, and applied the D-NEMD method with guidance by ASFO and the *DynaComm.py* code as advised by SO. MC set up and carried out equilibrium molecular dynamics simulations, thereafter conducting DF analysis and helping with the Figures. FM carried out molecular dynamics simulations with Anisotropic Thermal Diffusion. SO, ASFO, AJM, and GC provided guidance in producing the manuscript. All authors have written and reviewed parts of the manuscript and agree with the contents of this study.

## Acknowledgements

SAS wishes to thank Prof. Alemayehu Gorfe (Univesity of Texas Medical School – Houston) for providing an initial structure of membrane-bound K-Ras4B, Prof. Ana Vila Verde (University of Duisburg-Essen) for assistance with her inorganic phosphate forcefield parameters, and Prof. Aleksander Lyubartsev (Stockholm University) for assistance with the *Slipids* forcefield. Authors at the University of Pavia acknowledge support from the Italian *Ministero dell’Università e della Ricerca* (MUR) and the University itself through the program *Dipartimenti di Eccellenza 2023– 2027*.

## SYNOPSIS AND TOC

### Do you speak Allosteric?

Starting from unbiased MD simulations of the well-known oncotarget K-Ras4B, we assess performance of four different approaches in detecting its allosteric behavior. Each approach brings its own unique insights and shows consistency with experiment.

