## Supporting Information for "Decrypting the languages of allostery in membrane-bound K-Ras4B using four complementary *in silico* approaches"

#### Contents

CONTENTS S1

STARTING STRUCTURE FOR EQUILIBRIUM MD AND VIEW OF KEY K-RAS4B COMPLEXES S3

MD PREPRODUCTION DETAILS S4

**Minimization** S4

**100 ps *NVT* Equilibration** S4

**100 ps *NpT* Equilibration** S5

INPUT, MATRIX DERIVATION, AND FURTHER DETAILS ON SPM S5

**Alignment and Generation of Average Structure for Clustering** S5

**Clustering** S6

**Calculation of Average Distance and Correlation Matrices** S6

**Full Details on SPM Branches I-V** S6

FURTHER D-NEMD DETAILS S7

**Isolation of "Reactive" Frames from Equilibrium MD simulations** S7

**Reconstruction of GDP and $[H_2PO_4]^-$ from $H_2O$ and GTP** S9

**Testing whether $[H_2PO_4]^-$ Geometry is Preserved during D-NEMD** S12

NORMALIZATION OF PER-RESIDUE SCORES FOR THE FOUR METHODS S13

CRITICAL ASSESSMENT OF THE CHOSEN METHODS S13

BIBLIOGRAPHY S15

### Starting Structure for Equilibrium MD and View of Key K-Ras4B Complexes

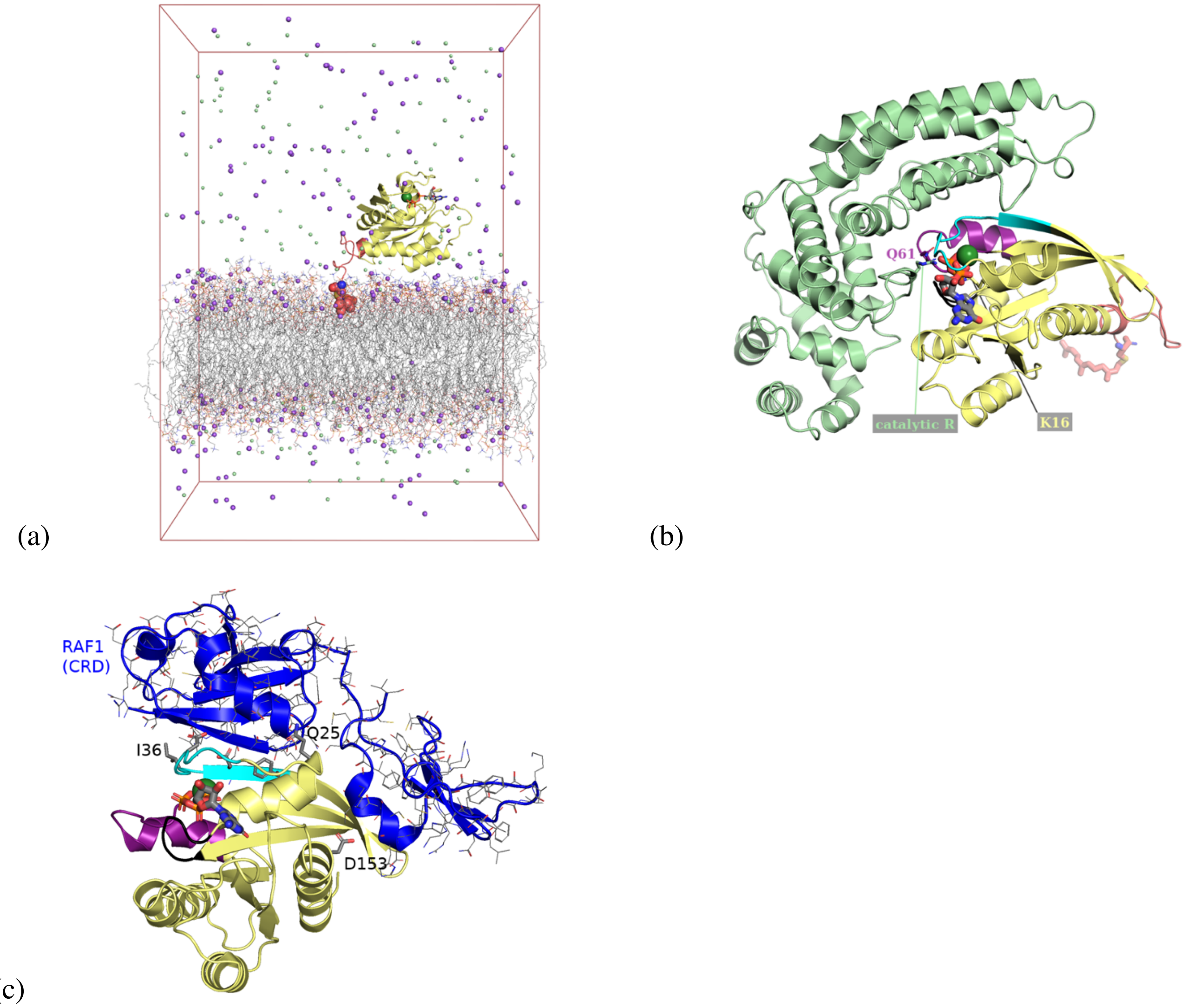


*Figure S1*. (a) Starting structure and periodic cell for simulations, set up as discussed in Computational Methods. The G domain (residues 2-166; yellow); GTP (sticks); and $Mg^{2+}$ (dark green sphere) are modeled after PDB ID: 6vjj.[1] The hypervariable region (HVR; residues 167-185; salmon) and the POPS/POPC phospholipid bilayer (lines) in which it is embedded through farnesylated Cys185 (C as large salmon spheres) are modeled from previous simulations.[2] Solute ions $Na^+$ and $Cl^-$ are rendered as purple and pale green spheres, respectively. (b) View from a different angle to reveal the location of the K-Ras4B—GAP interface (GAP in pale green; PDB ID: 1wq1),[3] showing all salient regions of the former's G-domain with the same color code as Figure 1. A catalytically crucial arginine administered by GAP (Arg789) is labeled and rendered as sticks and shown alongside Lys16 and Gln61; $Na^+$ and $Cl^-$ are omitted. (c) View from yet another angle to reveal the location of the K-Ras4B—RAF1 effector interface (cysteine-rich domain of RAF1; pale blue; residues rendered as lines; PDB ID: 6xi7).[1] Color code otherwise identical to Figure 1. Asp153 on helix α5 of K-Ras4B and other representative residues common to several effector interfaces are also rendered as sticks and/or labeled. $Na^+$ and $Cl^-$ are omitted. In all panels, H and solvent are omitted for clarity. Color code for explicit non-C atoms/ions: $Mg^{2+}$: dark green; Cl: pale green; Na: purple; P: orange; O: red; N: blue; S: yellow.

#### MD Preproduction Details

Run input files for all preproduction stages are provided electronically. Unless explicitly mentioned, parameters employed are either set to *GROMACS* defaults,[4] or identically to the MD production stage (see main text).

##### Minimization

Starting from the equilibrated, solvated, membrane-embedded K-Ras4B structure set up as discussed in the main text, all equilibrium MD replicas were begun by a round of steepest-descent structural minimization stopping once all atomic forces dropped below a 1000 kJ $mol^{-1}$ $nm^{-1}$ threshold to machine precision (usually taking about 2000 steps). No constraints were present at this stage, including on bonds containing hydrogens. In addition, the cutoff for the calculation of Lennard-Jones and Coulomb interactions was 8 Å rather than the 12 Å used at later stages. Beyond this limit, as usual, Coulomb interactions were computed in reciprocal space, with the *Particle Mesh Ewald* method[5] rather than in direct space, and Lennard-Jones interactions were set to zero (at this stage with no corrections).

##### 100 ps *NVT* Equilibration

Following minimization, all replicas were subjected to an initial 100 ps round of equilibration in the *NVT* ensemble ($T$ = 300 K): it was at this stage that atomic velocities for the entire system (a different set for each replica) were randomly assigned to match a Boltzmann distribution. Compared to the production stage, the timestep for the leap-frog integrator[6] was shorter, at 1 fs: to compensate, neighboring atom lists were made to update every 20 integrator steps rather than every 10. For this (and the next) equilibration stage, we imposed harmonic restraints ($k$ = 1000 kJ $mol^{-1}$ $Å^{-2}$ in all directions) on protein backbone heavy atoms; the $Mg^{2+}$ cation; all GTP atoms; all lipid heavy atoms.

Temperature enforcement conditions and couplings were identical to those used in the production stage. The cutoff for the computation of Lennard-Jones and Coulomb interactions was lengthened to

12 Å from this stage onwards; energy and pressure corrections to discrepancies arising from Lennard-Jones truncation were henceforth applied beyond this cutoff. Also from this stage onwards, *LINCS*[7] (non-water) and *SETTLE*[8] (water) constraints were introduced for all hydrogen-containing bonds.

##### 100 ps *NpT* Equilibration

After *NVT* equilibration, there followed a further 100 ps of equilibration, this time in the *NpT* ensemble ($T$ = 300 K; $p$ = 1 bar), with all previous atomic restraints retained. Contrarily to the production stage, constant pressure was in this case maintained by the Parrinello-Rahman barostat,[9] always with a time constant of 2 ps. Coupling to the barostat was still semi-isotropic, as described in the main text. Neighboring atoms continued to be searched for every 20 steps, but for this stage only, cutoffs for neighbor searching were temporarily shortened to 8 Å. All other conditions present in the previous equilibration stage were retained.

#### Input, Matrix Derivation, and Further Details on SPM

All input generation discussed in this subsection is carried out starting from our 5 $\mu$s metatrjectory. All steps are directly performed with the *cpptraj* tool,[10] whose input for each step we provide electronically.

##### Alignment and Generation of Average Structure for Clustering

Prior to clustering, our K-Ras4B metatrajectory is processed as follows:

- All frames are aligned to the very first metatrajectory frame (originally from replica 1), based on minimizing the root-mean-square deviation (RMSD) of Cα atoms. Cα atoms of HVR residues are included in this and subsequent alignments.
- The resulting Cα-aligned metatrajectory is used to derive an average structure ("average" command in *cpptraj*).[10]
- The Cα-aligned metatrajectory is realigned to Cα atoms of the average structure.

#### Clustering

Clustering is performed on every 50th frame of the realigned metatrajectory, or more specifically, after retaining every 25th frame (one every 50 ps), and sieving every second frame from this pruned metatrajectory to perform clustering. The algorithm used is the average-linkage hierarchical agglomerative approach as implemented in *cpptraj*.[10] The criterion on which to cluster was the RMSD of Cα atoms across the pruned metatrajectory, and the clustering run was made to stop once five clusters were found. The representative structure for the most populated cluster (lowest distance to centroid) was used for the final realignment of the (non-pruned) metatrajectory (see next subsection).

#### Calculation of Average Distance and Correlation Matrices

Alignment of each frame of the original metatrajectory to the representative structure of the most populated cluster was again carried out based on RMSD of Cα atoms, after which, the "matrix" command in *cpptraj*[10] is used twice in succession to produce both the distance matrix and the pairwise correlation matrix (see input file provided electronically for syntax; we also provide the matrices themselves). As *per* formulae provided in the main text, *DynaComm.py*[11] then uses these matrices directly (together with a chosen structure of the protein at hand) to work out which node-node edges should be represented in the pre-SPM graph (*i.e.*, those linking residues closer than 6 Å), and how heavy these edges should be.

#### Full Details on SPM Branches I-V

We here describe the paths of SPM branches I-V picking them up as they eradiate from the central allosteric hub. An overview of Branch V and its subbranch V.3 is provided in the main text.

Branch I (Figure 4; bottom) returns to Glu162 from Pro110 and conveys allosteric signals along the almost entirety of helix α5, incidentally avoiding the flexible Asp153 identified by DF analysis. Branch II (Figure 4; bottom) reverberates to sheet β6, through which it travels in its entirety before reaching Ser145 and Ala146 on the β6–α5 loop and therefore the binding site (guanine moiety), and further jumping across to Leu19 on helix α1. Branch III (Figure 4; bottom) terminates abruptly at Ala134 in the *C*-terminus of α4, after crossing to the α4–β6 loop. Branch IV (Figure 4; bottom) departs Pro110 and moves back towards the *N*-terminus along the α3–β5 loop, dissipating at Ser106.

As stated, Branch V branches off to β4 from the hub on β5, with a dual coupling to from Pro110 (β5) to Gly77 (β4) and from Met111 to Phe78 (β4); from there it further branches out into three subbranches V.1, V.2, and V.3. V.1 crosses almost the entirety of sheet β1 and V.2 just reaches the *C*-terminal Arg73 on α2, which is just outside of switch II. V.3 is discussed in the main text.

#### Further D-NEMD Details

##### Isolation of "Reactive" Frames from Equilibrium MD simulations

Our MD metatrajectory comprises a total of 100000 frames saved with stored atomic velocities. 20 of these are automatically unviable for D-NEMD, as they are saved at the very end of their respective equilibrium replica: D-NEMD simulations begun from these frames would have no equilibrium counterpart against which to monitor Cα deviation. Of the remaining 99980 frames (4999 per replica), we omit the first 100 frames of each replica (further lowering the total of viable frames down to 4899), and we furthermore only begin D-NEMD simulations from those 71706 frames that, based chemical considerations from the previous QM/MM study by Calixto and coworkers,[12] meet certain criteria. Frames recognized as "reactive" come from across all 20 MD replicas and are tabulated Table S1.

Table S1. Summary of the total frames (windows) on which D-NEMD simulations were conducted, and those that survived for their whole 50 ps duration.

| **Replica** | **Viable frames** | **"reactive" frames without considering Q61 closure** | **Surviving after D-NEMD** | **Loss** | **"reactive" frames considering Q61 closure** | **Surviving after D-NEMD** | **Loss** |
|---|---|---|---|---|---|---|---|
| 1 | 4899 | 3743 | 2921 | 22.0% | 932 | 628 | 32.6% |
| 2 | 4899 | 3716 | 2973 | 20.0% | 1938 | 1518 | 21.7% |
| 3 | 4899 | 3720 | 2883 | 22.5% | 920 | 588 | 36.1% |
| 4 | 4899 | 3689 | 2781 | 24.6% | 1006 | 601 | 40.3% |
| 5 | 4899 | 3939 | 3464 | 12.1% | 3231 | 2888 | 10.6% |
| 6 | 4899 | 4296 | 3539 | 17.6% | 2564 | 2146 | 16.3% |
| 7 | 4899 | 3745 | 3207 | 14.4% | 2748 | 2394 | 12.9% |
| 8 | 4899 | 3580 | 2845 | 20.5% | 1518 | 1205 | 20.6% |
| 9 | 4899 | 4319 | 3151 | 27.0% | 1079 | 575 | 46.7% |
| 10 | 4899 | 4112 | 3212 | 21.9% | 997 | 635 | 36.3% |
| 11 | 4899 | 1898 | 1522 | 19.8% | 611 | 478 | 21.8% |
| 12 | 4899 | 2991 | 2525 | 15.6% | 611 | 500 | 18.2% |
| 13 | 4899 | 2980 | 2512 | 15.7% | 2124 | 1848 | 13.0% |
| 14 | 4899 | 2956 | 2550 | 13.7% | 1440 | 1249 | 13.3% |
| 15 | 4899 | 4099 | 3074 | 25.0% | 1382 | 947 | 31.5% |
| 16 | 4899 | 3116 | 2587 | 17.0% | 1683 | 1425 | 15.3% |
| 17 | 4899 | 3597 | 3056 | 15.0% | 2156 | 1855 | 14.0% |
| 18 | 4899 | 3346 | 2916 | 12.9% | 638 | 561 | 12.1% |
| 19 | 4899 | 4089 | 3059 | 25.2% | 1029 | 637 | 38.1% |
| 20 | 4899 | 3775 | 3046 | 19.3% | 793 | 531 | 33.0% |
| metatrajectory (Total) | 97980 | 71706 | 57823 | 19.4% | 29400 | 23209 | 21.1% |

Criteria that should simultaneously be met by an equilibrium MD frame to qualify as "reactive" are as follows:

1. That there exists a water molecule (the nucleophile), whose oxygen should be:
    a. no farther than 3.70 Å from the Pγ atom in GTP; and
    b. poised to attack GTP with an angle of >145° with respect to the Pγ–O3β bond.
2. That at least one of the three Hζ hydrogens in Lys16—which is important[12] to contrast the accumulation of negative charge in the transition state during GTP hydrolysis—is no farther than 2.19 Å from at least one of the Oγ (terminal) atoms of GTP.

In addition, optionally (see main text; *Results and Discussion*), we opted to recalculate D-NEMD statistics with a more stringent (and chemically plausible) criterion to determine reactive poses:

3. That the Cδ atom in Gln61, which under the right conformational conditions helps position the nucleophile for attack,[12] is no farther than 4.45 Å from one of the two hydrogens in the nucleophilic water molecule.

Since, in the absence of a GAP protein, Gln61 is conformationally freer, the inclusion of the third criterion greatly reduces the number of reactive poses (*cf.* Table S1), while not affecting the final results (data not shown). All of the (rather lax) thresholds in criteria 1-3 above were decided based on radial distribution functions involving the atoms in question (or histogram, in the case of the nucleophilic attack angle), choosing them to include the entire first peak (data not shown).

#### Reconstruction of GDP and $[H_2PO_4]^-$ from $H_2O$ and GTP

Our main objective when automatically introducing the perturbation at the start of each D-NEMD window was to instantaneously reproduce the $GTP^{4-} + H_2O \rightarrow GDP^{3-} + [H_2PO_4]^-$ reaction as closely as possible by changing the positions of only 4 atoms: Pγ in GTP; and the three atoms composing the nucleophilic water molecule. In this reaction, one of the γ-oxygens of GTP is acting as the base that deprotonates the nucleophilic water.[12] The beginning and outcome of a typical reconstruction are illustrated in Figure S2.

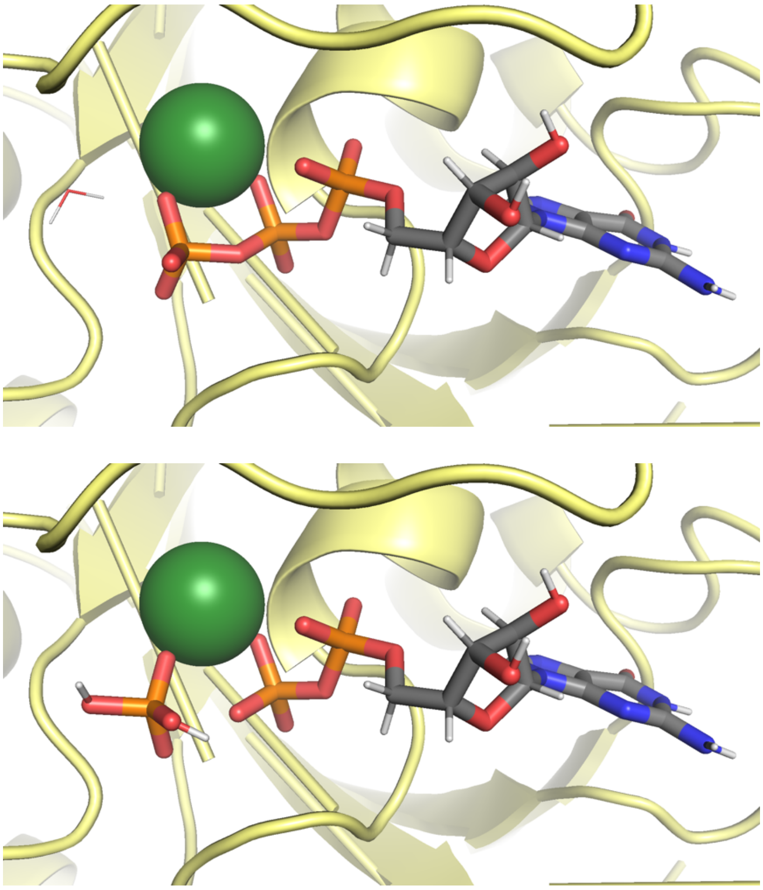

*Figure S2*. Example of the transformation (perturbation) performed at the beginning of each D-NEMD run on the corresponding "parent" pose issuing from equilibrium MD, whereby $GTP^{4-}$ and a nucleophilic water (top) are converted to $GDP^{3-}$ + $[H_2PO_4]^-$ (bottom) by only moving four atoms. The resulting $[H_2PO_4]^-$ (bottom) is constructed at equilibrium geometry. Color codes are identical to Figure 1b, except that the salient G-domain regions are left in yellow. All solvent molecules apart from the nucleophilic water have been omitted.

Using an in-house script, and relying on the *NumPy* module in *Python*, the steps taken to enact hydrolysis with minimal change were as follows:

1. Umbrella (pyramidal) inversion of GTP's terminal (γ-)phosphate, by shifting the Pγ atom as follows:
    a. Identification of the plane containing the three Oγ atoms in GTP (O1γ, O2γ, O3γ; leftmost oxygen GTP oxygen atoms in Figure S2 top).
    b. Identification of the normal to this plane.
    c. Calculation of the distance of Pγ from this plane as the dot product between the plane's normal and the Pγ → O1γ vector (*i.e.*, its projection onto the plane).
    d. Displacement of Pγ along the Pγ → O3β vector (*i.e.*, in-axis with the Pγ–O3β bond to be cleaved) by twice the distance between Pγ and the Oγ plane, so that Pγ

"reappears" on the other side of the plane but at the same distance, effectively severing the Pγ – O3β bond and producing a GDP molecule.

2. Displacement of the oxygen in the nucleophilic water molecule ($O_{wat}$) to create a new, *post*-hydrolysis $O_{wat}$–Pγ bond, of the 1.625 Å length prescribed by $[H_2PO_4]^-$ forcefield parameters,[13] again in-axis with the former Pγ–O3β bond. This effectively tears the nucleophilic water apart.
3. Displacement of one of the hydrogens on nucleophilic water ($H_{wat1}$) to "catch up" with $O_{wat}$ shifted in step 2, at the prescribed[13] geometry for $[H_2PO_4]^-$. This means: a $O_{wat}$–$H_{wat1}$ bond length of 0.974 Å, a $H_{wat1}$–$O_{wat}$–Pγ angle size of 110.14°, and a $H_{wat1}$–$O_{wat}$–Pγ–$O\gamma_{base}$ dihedral of 180°, where Oγ is the γ-phosphate oxygen chosen to act as a base. Typically, this is the oxygen that is farthest from Lys16 and $Mg^{2+}$. Displacement is achieved as follows:
   a. Identification of the plane containing $H_{wat1}$ in its old position, $O_{wat}$ in its new position from step 2, and Pγ in its new position from step 1.
   b. Identification of the unit normal to this plane $\vec{\mathbf{k}}$, passing through atom $O_{wat}$ in its new position, given by the normalized cross product between vectors $O_{wat} \rightarrow$ Pγ and $O_{wat} \rightarrow H_{wat1}$.
   c. Measurement of the old $H_{wat1}$–$O_{wat}$–Pγ angle, from $O_{wat} \rightarrow$ Pγ and $O_{wat} \rightarrow H_{wat1}$.
   d. Application of Rodrigues' formula to rotate the $O_{wat} \rightarrow H_{wat1}$ vector about $O_{wat}$ so that the $H_{wat1}$–$O_{wat}$–Pγ angle attains the prescribed 110.14° after rotating $\theta$ degrees from the old angle identified in 3c. Rodrigues' formula is as follows:

$$\overrightarrow{OH}_{new} = \overrightarrow{OH}_{old} \cos\theta + \left(\vec{\mathbf{k}} \times \overrightarrow{OH}_{old}\right) \sin\theta + \vec{\mathbf{k}}(\vec{\mathbf{k}} \cdot \overrightarrow{OH}_{old})(1 - \cos\theta)$$

   where $\vec{\mathbf{k}}$ was defined in 3b, and $\overrightarrow{OH}_{old}$ and $\overrightarrow{OH}_{new}$ denote the $O_{wat} \rightarrow H_{wat1}$ before and after rotation, respectively. This operation only yields the correct $H_{wat1}$–$O_{wat}$–Pγ angle; lengths and dihedrals are fixed in the next substeps.

e. Shortening the new $O_{wat} \rightarrow H_{wat1}$ vector to its desired equilibrium length of 0.974 Å[13] (by normalizing it to a unit vector and multiplying the latter by 0.974 Å).

f. Measurement of the old dihedral $H_{wat1}$–$O_{wat}$–Pγ–$O\gamma_{base}$ dihedral. Requires finding the normal vectors to the planes $H_{wat1}$–$O_{wat}$–Pγ and $O_{wat}$–Pγ–$O\gamma_{base}$, and taking the arccosine of their dot product.

g. Rotation of the $H_{wat1}$–$O_{wat}$–Pγ–$O\gamma_{base}$ dihedral about the $O_{wat}$–Pγ bond, from the old dihedral angle to the prescribed 180°,[13] using the same formula in 3d.

h. Establishment of the final "new" coordinates of $H_{wat1}$ after the rotation.

4. Displacement of the remaining hydrogen on nucleophilic water ($H_{wat2}$)—now the only one remaining in its old position—to $O\gamma_{base}$, again at the prescribed[13] geometry for $[H_2PO_4]^-$. This is done following precisely the same substeps as in 3, except that the bond to regulate is now $O\gamma_{base}$–$H_{wat2}$; the angle is now $H_{wat2}$–$O\gamma_{base}$–Pγ; and the dihedral is now $H_{wat2}$–$O\gamma_{base}$–Pγ–$O_{wat}$.

The $[H_2PO_4]^-$ anion resulting from this operation—at equilibrium geometry[13]—is shown in Figure S2 bottom. Note that positions of O1γ, O2γ, O3γ, and O3β have remained unchanged, and only four atoms have moved.

#### Testing whether $[H_2PO_4]^-$ Geometry is Preserved during D-NEMD

To check for any unphysical breakups of the $[H_2PO_4]^-$ anion during D-NEMD equilibration in a particular window, using an in-house script relying on the *Biopython* package[14] for quicker access to vectors we automatically perform the following checks:

- That none of the $[H_2PO_4]^-$ bonds have elongated or shortened by more than 20% compared to equilibrium geometry.
- That none of the $[H_2PO_4]^-$ angles have opened or closed by more than 20% compared to equilibrium geometry.

- That $[H_2PO_4]^-$ dihedrals are no more than 35% of the way towards their maxima.

#### Normalization of Per-residue Scores for the Four Methods

The per-residue raw score $S_{raw}$ chosen for each allosteric language and plotted in Figure 7 is rescaled/normalized so that the score $S_{norm,i}$ for the $i^{th}$ residue will always fall between 0 and 1. Normalization is enacted through the following formula:

$$S_{norm,i} = \frac{S_{raw,i} - S_{raw,min}}{S_{raw,max} - S_{raw,min}}$$

wherein $S_{raw,min}$ and $S_{raw,max}$ denote, respectively, the minimum and maximum raw scores detected for a particular method over the 168 residues considered. Essentially, therefore, the normalized score $S_{norm,i}$ measures how close the raw score for the $i^{th}$ residue $S_{raw,i}$ is to either $S_{raw,min}$ or $S_{raw,max}$ on a scale from 0 to 1, *i.e.*, it has a value of 0 for the lowest-scoring residue ($S_{raw,i} = S_{raw,min}$) and 1 for the highest-scoring residue ($S_{raw,i} = S_{raw,max}$), and falls in between these extremes for remaining residues.

#### Critical Assessment of the Chosen Methods

To recap some of the considerations specifically applying to our own choice of methods, SPM and DF clearly have the greatest advantage in terms of computational cost since, as we recounted in the main text, they can extract allosteric information from just one series of unbiased MD simulations (D-NEMD and ATD require additional integrative simulations). "Equilibrium" languages SPM and DF both have their own unique strengths and weaknesses. For example, DF allows a unique two-dimensional breakdown of allosteric cross-talk signals at equilibrium (the DF score matrix; Figure 3, bottom), which as we have seen is useful, since not all allosterically active regions dialogue with each other with the same intensity. On the other hand, 'flattening' the DF matrix leads to an average 1D score that can be projected onto a structure under examination (*e.g.* Figure 3, top/middle), but loses a considerable amount of spatial information, *i.e.*, one only retains information on tendentially rigid *vs.* tendentially uncoordinated residues. The SPM approach provides an elegant way to circumvent this problem: while still starting from the multidimensional information about how each residue talks to every other residue, the allosteric maps that are produced (*e.g.*, Figure 4) conveniently condense the information into a linear 'shortest path', which obviously not only retains spatial information about the "flow" of allosteric signals, but also quantitative information about their changing intensity.

One *caveat* of SPM, however, is that being based on average distances and covariances (*cf. Computational Methods*) it requires that they be measured after structural alignment to a sufficiently representative reference structure: this needs to be derived carefully, as the choice of clustering and alignment can somewhat impact results, particularly in path sections close to the 0.3 'shortness' threshold. The *caveat* does not apply to DF analysis, which only monitors intra-frame Cα–Cα distances regardless of orientation, and therefore only requires that a molecule in a particular frame is whole. Both methods are nonetheless also still somewhat limited by the conformational space sampled along the reference MD simulations.

While perturbative methods, such as D-NEMD and ATD, require extra simulations and therefore come at an increased computational cost, it is a price worth paying given the level of allosteric insight that they are able to provide: with the notable advantage of not requiring to model a conformational change, both these languages can offer an accurate indication of what happens allosterically upon a specific perturbation, when only a part of the equilibrium pathways will be exalted over others. Though it may be argued that the perturbation introduced by these methods is a form of bias, this is not strictly true, insofar as they do not accelerate a particular conformational transition but, still retaining equilibrium MD simulations as their standard, they merely try and detect which areas of the system vary the most once the perturbation is introduced. Conveniently, the perturbation does not have to make strict physicochemical sense, as is the case with both the supercooling and decoupled heating in ATD and the instantaneous GTP hydrolysis in D-NEMD.

A particularly elegant aspect of D-NEMD (Figure 2) is that the non-equilibrium simulations are initiated instantaneously from equilibrium ones, with the same set of atomic velocities, and deviation is monitored systematically at synchronous intervals. This essentially reduces the noise that may arise from inherently flexible regions, because flexible regions that are not affected by the perturbation under examination will, at short timeframes, in principle deviate by identical amounts both at equilibrium and at non-equilibrium; the only appreciable source of deviation between the two situations is thus bound to be ascribable to the perturbation itself.

ATD, on the other hand, is less capable of eliminating noise, since deviations in each window are monitored from a fixed time reference (*i.e.*, the start of heating for that window): this means that it is impossible to entirely separate deviations that simply happen because of flexibility, and those that happen because they have received an allosteric impulse from the perturbation, and it is the reason why the HVR had to be excluded from the $S_{\mathrm{norm}}$ plots in Figure 7. The attractivity of ATD lies in the fact that it can act as a sort of 2D "allosteric NMR", selectively overstimulating the contribution of one particular residue, sets of residues, or as in our case, of course, the nucleotide itself.
